# Recombinase polymerase amplification: characterization and mitigation of undescribed multimeric artefacts

**DOI:** 10.64898/2026.08.21.741777

**Authors:** Liesl De Keyzer, Koen Deserranno, Sonja Skevin, David Van Hoofstat, Dieter Deforce, Filip Van Nieuwerburgh

## Abstract

Recombinase polymerase amplification (RPA) enables rapid nucleic acid testing in low-resource environments, but poorly characterized byproducts can compromise assay specificity and cause false-positive results. Here, we amplified the thirteen original CODIS core loci and Amelogenin to characterize recurrent RPA artefacts and establish conditions that reduce their formation. First, RPA products were analyzed for two reference samples by Oxford Nanopore Technologies sequencing. This revealed two distinct classes of multimeric products: primer multimers and amplicon multimers, consisting of repeated primer or amplicon sequences, respectively. Individual artefacts contained up to 281 primer copies or 22 amplicon copies, demonstrating the extensive range of these products. Next, we performed an optimization study to evaluate the effects of reaction temperature and reagent concentrations at two representative loci, D3S1358 and D5S818. Among the conditions tested, temperature had the most pronounced effect. Reducing the temperature from 42°C to 34°C increased the relative target amplicon fraction from 15% to 83% for D3S1358 and from 84% to 98% for D5S818, while maintaining or increasing absolute target concentration. Lower primer concentrations and higher T4 UvsX concentrations also reduced multimer formation, although lower primer concentrations reduced target yield and caused allelic dropout. Finally, amplification at 34°C was evaluated across all fourteen loci by sequencing. Relative to 42°C, the target read fraction increased by more than 5 percentage points for 7/14 loci in one reference sample and 9/14 loci in the other, with the largest improvements at multimer-prone loci. These findings identify multimers as an important class of RPA artefacts and establish reaction temperature and T4 UvsX concentration as promising conditions to improve RPA specificity.

## Introduction

Among nucleic acid amplification techniques, polymerase chain reaction (PCR), published by Mullis et al. in 1986 (1), is still regarded the gold standard. However, PCR requires precise thermal cycling and therefore depends on calibrated, bulky instrumentation, which limits its use in low-resource environments (2,3). Over the past two decades, recombinase polymerase amplification (RPA) has emerged as an important isothermal alternative, with reports in more than 2000 publications on PubMed since its invention by Piepenburg et al. in 2006 (4). RPA relies on a recombinase enzyme to facilitate primer invasion into the DNA duplex, and a strand-displacing polymerase for primer elongation (5). As amplification proceeds without a thermal denaturation step, RPA can be performed at a low and constant temperature of 37-42°C, resulting in exponential amplification in 5-20 minutes (6). These characteristics make RPA particularly suitable for applications outside laboratory settings where conventional thermocycling equipment is unavailable or impractical, for instance in point-of-care genetic screening (7), biosensing (8), and rapid diagnostics (9).

Despite its advantages, RPA is prone to the formation of nonspecific reaction products, which are claimed to be formed by unintended primer interactions due to recombinase’s limited capacity to discriminate between mismatches (10–12). These artefacts may lead to false-positive results and reduce overall reaction specificity and sensitivity. In addition, off-target amplification may interfere with the preparative synthesis of target sequences intended for downstream analyses (13). Despite its significance, nonspecific product formation in RPA has not been extensively studied, with only a limited number of publications specifically addressing this issue (12–15). Although the presence of these reaction artefacts is frequently observed as distinct bands on agarose gels (16–19), their structural composition and the reaction dynamics underlying their formation remain poorly characterized.

In the field of forensic DNA identification, access to real-time information at crime scenes can be highly valuable for the guidance of police investigations during the critical early phase of a case (20). However, DNA analysis turnaround times often remain longer than desired, ranging from several weeks to months, which can delay decision-making across the criminal justice chain (21). Consequently, there is a growing interest in bringing forensic DNA analysis closer to the crime scene, creating a need for portable, straightforward, and robust analytical platforms (22). RPA is well aligned with this objective due to its compatibility with lab-on-a-chip devices and simple readout formats (7). However, forensic samples often contain low amounts of DNA and may be degraded or present as mixtures, requiring assays with high sensitivity and specificity (23). In our previous studies, we demonstrated that RPA can be used for forensic short tandem repeat (STR) (19) and InDel genotyping (6), although specificity-related challenges were observed for both markers.

In this study, we amplified the 13 original forensic CODIS core loci and Amelogenin with RPA, to investigate both the structural composition of RPA artefacts and the effect of reaction conditions on their formation. A dual analytic approach was applied. First, pooled singleplex RPA products were sequenced using Oxford Nanopore Technologies (ONT), revealing two distinct types of RPA artefacts, hereafter referred to as primer multimers and amplicon multimers. These multimers were characterized by repeated primers or amplicons arranged in a head-to-tail fashion. To the best of our knowledge, these artefacts have not yet been described in literature. Second, the influence of several key reaction conditions on RPA artefact formation was evaluated using Agilent’s Fragment Analyzer. Among the tested parameters, reaction temperature had the strongest effect, with lower temperatures reducing multimer formation. Overall, these findings provide insight into the structure and formation of nonspecific RPA products and may support broader assay optimization in applications where RPA specificity is critical.

## 2. Materials and methods

### 2.1. Sample collection and purification

This study was performed using two commercially available reference DNA samples, 9947A and 9948 (OriGene, Rockville, Maryland, USA), and one blood sample obtained from a healthy volunteer. Ethical approval was granted by the ethical review board of Ghent University Hospital (approval No. BC-05557), and participants provided written informed consent prior to blood donation. Blood was collected by finger prick using a 21 G Minicollect^®^ Lancelino safety lancet with a penetration depth of 2.4 mm and transferred into K3E K3EDTA Minicollect^®^ collection tubes (Greiner Bio-One, Kremsmünster, Austria). The DNA was extracted using the DNeasy^®^ Blood and Tissue kit (Qiagen, Hilden, Germany) following the manufacturer’s protocol. Reference DNA samples were used without additional preparations. All samples were diluted in nuclease-free water to a final concentration of 1 ng/µL for downstream analysis.

### 2.2. Benchmark genotyping

All included samples were genotyped for the 13 original CODIS core loci and Amelogenin by Eurofins Forensics Belgium NV (Bruges, Belgium) using capillary electrophoresis (CE). First, 1 ng DNA was amplified for 28 PCR cycles using the AmpFlSTR^®^ Identifiler^®^ Plus PCR Amplification Kit (Thermo Fisher Scientific, Waltham, MA, USA) according to the manufacturer’s instructions. Next, 1 µL of amplified product was loaded on the ABI3500xl Genetic Analyzer (Thermo Fisher Scientific, Waltham, MA, USA). The resulting electropherograms were analyzed using GeneMapper ID-X 1.2 software (Thermo Fisher Scientific, Waltham, MA, USA). The true genotypes of all samples are provided in Supplementary Table S1.

### 2.3. Recombinase polymerase amplification

RPA reactions were performed using the Lyo-Ready RPA kit (Invitrogen, Waltham, MA, USA), according to the manufacturer’s instructions. For standard reactions, a master mix was prepared to achieve final concentrations of 1 × Reaction Buffer, 1.1 mM dNTPs, 0.06 mg/mL T4 UvsX protein, 0.03 mg/mL T4 UvsY protein, 0.4 mg/mL T4 Gene 32 protein and 0.15 U/µL Bst DNA polymerase. Next, 1 ng of DNA was added to the mastermix, followed by the RPA primers at a final concentration of 0.3 µM. Reactions were initiated by the addition of MgCl_2_ at a final concentration of 14 mM, bringing the total reaction volume to 20 µL. All samples were mixed and incubated at 42°C for 40 minutes in a SimpliAmp Thermal Cycler (Life Technologies, Waltham, MA, USA). After 4 min of incubation, samples were briefly mixed by inversion before continuing amplification for the remainder of the 40 minutes run. Post-amplification, samples were purified using AMPure XP beads (Beckman Coulter Life Sciences, Indianapolis, IN, USA) at a sample-to-bead ratio of 1:1.8, according to the manufacturer’s instructions. Final elution was performed in 15 µL of nuclease-free water. The RPA primer sequences and locus information are given in Supplementary Table S2.

### 2.4. Oxford Nanopore sequencing

To identify and characterize the RPA artefacts, reference samples 9947A and 9948 were amplified in singleplex for the 13 original CODIS core loci and Amelogenin. RPA amplification was performed at two reaction temperatures, 42°C and 34°C, while keeping all other reaction conditions unchanged as described in Section 2.3. The resulting singleplex RPA products were quantified using the Qubit dsDNA High Sensitivity Assay kit and a Qubit 4.0 Fluorometer (Invitrogen, Waltham, MA, USA), according to the manufacturer’s protocol. Following quantification, the singleplex reactions were pooled equimolarly per sample. This resulted in an average input for library preparation of 356.59 ng for products amplified at 42 °C and 63.78 ng for products amplified at 34 °C. Libraries were prepared using the SQK-NBD114.24 kit, according to the manufacturer’s instructions (Oxford Nanopore Technology, Oxford, UK). Briefly, end preparation was performed using the NEBNext End Repair/dA-Tailing Module (NEB, Ipswich, MA, USA), followed by purification with a 1.8 × volume of AMPure XP beads (Beckman Coulter, High Wycombe, UK). Next, native barcode ligation was carried out using the NEB Blunt/TA Ligase Master Mix (NEB, Ipswich, MA, USA), followed by a 0.7 × AMPure XP bead clean-up. Finally, native adapter ligation was performed using the NEBNext Quick Ligation module (NEB, Ipswich, MA, USA), followed by another 0.7 × AMPure XP bead clean-up. The prepared libraries were quantified using the Qubit dsDNA High Sensitivity Assay kit (Invitrogen, Waltham, MA, USA). A total of 20 fmol DNA was loaded onto a PromethION R10.4.1 flow cell, according to the manufacturer’s instructions. Sequencing was performed on a PromethION 24 device for 72 h to maximize data output. Figure 1A gives an overview of the sequencing workflow.

**Figure 1:**
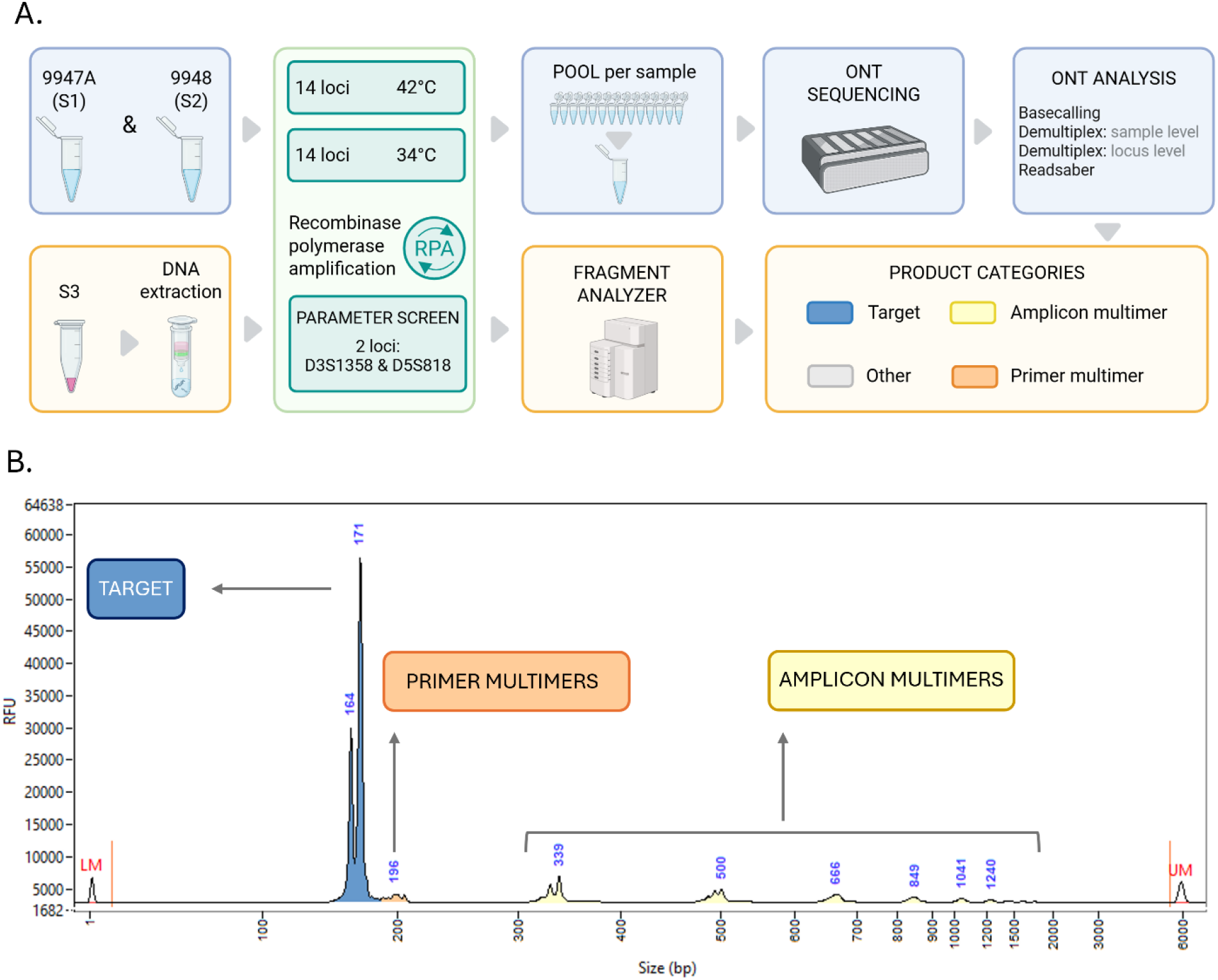
**A)** Overview of the sample preparation workflow. Sample 1 and 2 (reference DNA samples 9947A and 9948), were initially amplified for 14 loci in singleplex at 42°C, pooled per sample, and analyzed by ONT sequencing to identify and characterize RPA artefacts. Sample 3 was subsequently amplified for two selected loci, D3S1358 and D5S818, to evaluate the effect of reaction conditions on artefact formation. As amplification at 34°C reduced the artefact proportion for those two loci, sample 1 and 2 were sequenced again after amplification at 34°C to assess whether these findings could be extended to other CODIS core loci. **B)** Example of Fragment Analyzer peak annotation for locus D5S818. Target peak(s) were first annotated based on the expected amplicon length. Peaks larger than the target peak were annotated as primer multimers when their size increased by a multiple of the primer length, and as amplicon multimers when their size increased by a multiple of the expected amplicon length.

### 2.5. Sequencing data analysis

ONT sequencing reads were basecalled, trimmed, and demultiplexed using the dna_r10.4.1_e8.2_400bps_sup@v5.2.0 model of Dorado v1.1.0 (Oxford Nanopore Technologies, 2025), generating sample-specific FASTQ files. Reads with an average Q-score below 10 were discarded. Next, the sample-level FASTQ files were separated into 14 locus-specific FASTQ files based on the RPA primers using a custom Python script. Reads containing primer sequences from multiple loci were rejected to prevent incorrect locus assignment, accounting for 3.41 % of reads for 9947A and 1.45 % for 9948. The locus-specific FASTQ files were then used as input for annotation with Readsaber v0.1.1 (https://github.com/derijkp/readsaber) (Figure 1A). Briefly, Readsaber was used to annotate structural read elements, including native barcodes (NB), native adapters (NA), and primers. Target sequences were annotated with Readsaber by mapping the reads against a reference library containing all known STR alleles present in more than 1% of the world population, as collected from NIST STRbase (24). For each locus, Readsaber generated a summary figure displaying the most common annotation patterns. Based on the observed patterns, reads were classified as true amplicons, primer multimers, amplicon multimers, or other RPA artefacts. True amplicons were defined as reads solely containing the expected target amplicon sequence, flanked by the forward and reverse primer in the correct orientation. Primer multimers were defined as reads containing at least two primers arranged in a head-to-tail fashion. These products included both repeats of the same primers (e.g. forward primer – forward primer) and combinations of different primers (e.g. forward primer – reverse primer), provided that the primers had the same orientation and immediately followed each other. Two adjacent primers in opposite orientations were considered standard primer dimers instead of primer multimers, and were classified as ‘other’ RPA artefacts. Amplicon multimers were defined as reads containing at least two amplicons, potentially interspersed by other sequences, where amplicons were defined as target sequences flanked by one forward and one reverse primer in the correct orientation. Reads that did not match any of these patterns were classified as ‘other’ RPA products. STR genotyping was performed on the locus-specific FASTQ files. Reads were subjected to a custom filtering based on the alignment score (AS), as previously described by our group (19). Briefly, reads were first mapped to the library of STR alleles with bwa mem (v0.7.17), due to the short nature of the amplicons (25). Next, all reads with an AS above 90% of the read span were retained. Reads mapping to each allele were counted, and the allele with the highest coverage was called as present. A second allele was called when its read count exceeded 50% of that of the most abundant allele.

### 2.6. Optimization of reaction conditions

The influence of variable reaction parameters on RPA artefact formation was evaluated using sample 3 (Figure 1A). Based on the sequencing results for samples 9947A and 9948, two representative loci were selected for optimization: D3S1358, characterized by primer multimers, and D5S818, characterized by amplicon multimers. Sample 3 was subsequently amplified at both loci as explained in Section 2.3, while varying individual reaction parameters in a one-factor-at-a-time approach. An overview of the parameters and their test ranges is provided in Table 1. The resulting RPA products were analyzed using the 5200 Fragment Analyzer (Agilent, Santa Clara, CA, USA), by loading 2 µL of a 1:10 dilution according to the manufacturer’s instructions.

**Table 1.** RPA reaction parameters and test ranges.

| Reaction parameter | Range |
| --- | --- |
| Temperature ($^{\circ}$C) | 34 – 35 – 36 – 37 – 38 – 39 – 40 – 41 – 42 – 43 – 44 – 45 |
| DNA input (ng) | 0 - 0.25 - 0.5 – 1.0 - 2.5 – 5.0 – 7.5 – 10.0 |
| [Primer] (μM) | 0 - 0.1 - 0.2 - 0.3 - 0.4 - 0.5 - 0.6 |
| [dNTP] (mM) | 0.5 - 0.8 - 1.1 - 1.8 - 2.4 – 3.0 |
| [MgCl<sub>2</sub>] (mM) | 7.0 – 10.0 – 14.0 – 18.0 – 21.0 |
| [T4 UvsX] (mg/mL) | 0.015 - 0.03 - 0.06 - 0.09 - 0.12 - 0.15 |
| [T4 UvsY] (mg/mL) | 0.025 - 0.03 - 0.035 - 0.04 - 0.045 - 0.05 |
| [T4 GP32] (mg/mL) | 0.4 - 0.45 - 0.5 - 0.6 - 0.75 - 0.9 |
| [Bst DNA polymerase] (U/μL) | 0.015 - 0.03 - 0.06 - 0.09 - 0.12 - 0.15 |

### 2.7. Fragment analyzer results

Analysis of the Fragment Analyzer profiles was performed using the ProSize Data Analysis Software (Agilent, Santa Clara, CA, USA), as illustrated with an example in Figure 1B. First, the highest peak within the locus’ expected length range was annotated as the target peak. Samples were considered heterozygous if the second highest peak within that range reached at least 1/3 of the fluorescence intensity of the highest peak, in which case the second highest peak was also annotated as a target peak. Genotypes were considered correct when the lengths of the annotated target peaks matched the benchmark genotype lengths, taking into account the reported 5% deviation for sizing accuracy (26). Genotypes were classified as incorrect when allelic dropouts or drop-ins were observed, where allelic dropout was defined as the failure to detect one allele in a heterozygous sample, resulting in its incorrect classification as homozygous. Allelic drop-in was defined as the appearance of a spurious peak that did not contribute to a true allele. Artefact peaks were classified according to their size difference relative to the target peak. Peaks separated by a multiple of the length of one primer were defined as primer multimers. Depending on the locus, this corresponded to an additional 24–36 bp. Peaks separated by a multiple of the length of one amplicon were classified as amplicon multimers, which corresponded with 112-385 bp depending on the locus. Peaks that could not be classified under any of the previous definitions, or peaks shorter than the target peak, were grouped as ‘other’ and included all undefined RPA artefacts. Relative target, primer multimer, and amplicon multimer concentrations (%) were calculated by adding up the concentrations of all peaks assigned to each category, divided by the total concentration, and multiplied by 100. The total concentration (ng/µL) was defined by ProSize as the total concentration of all detected peaks, excluding markers, and regardless of whether peaks were annotated or not (27). Absolute target concentrations (ng/µL) were calculated by adding up the concentrations of all target peaks and multiplying it by 10 to correct for the initial 1:10 dilution.

## 3. Results

### 3.1. Identification of two RPA artefacts

ONT sequencing of two pooled singleplex samples, 9947A and 9948, revealed two distinct RPA artefacts, hereafter referred to as primer multimers and amplicon multimers. Figure 2A shows the Readsaber summary plots of two loci, D3S1358 and D5S818, representing examples of primer multimers and amplicon multimers, respectively. The remaining Readsaber plots are shown in Supplementary Figure S1, and an overview of the RPA products across all loci is shown in Figure 2B. Across the 14 analyzed loci, primer multimers were most prominent for D3S1358 (49.81% for 9947A and 11.25% for 9948), TPOX (44.84% for 9947A and 18.41% for 9948), and vWA (47.38% for 9947A and 21.81% for 9948). Amplicon multimers were mainly observed for D3S1358 (5.85% for 9947A and 7.90% for 9948), D5S818 (14.07% for 9947A and 8.47% for 9948), and TH01 (8.05% for 9947A and 12.95% for 9948). Multimers typically generated extensively long reads. For example, the longest primer multimer was seen for Amelogenin in sample 9948, containing 281 primers. The longest amplicon multimer was seen for D5S818 in sample 9947A and contained 22 times the target sequence (Supplementary Figure S2).

**Figure 2:**
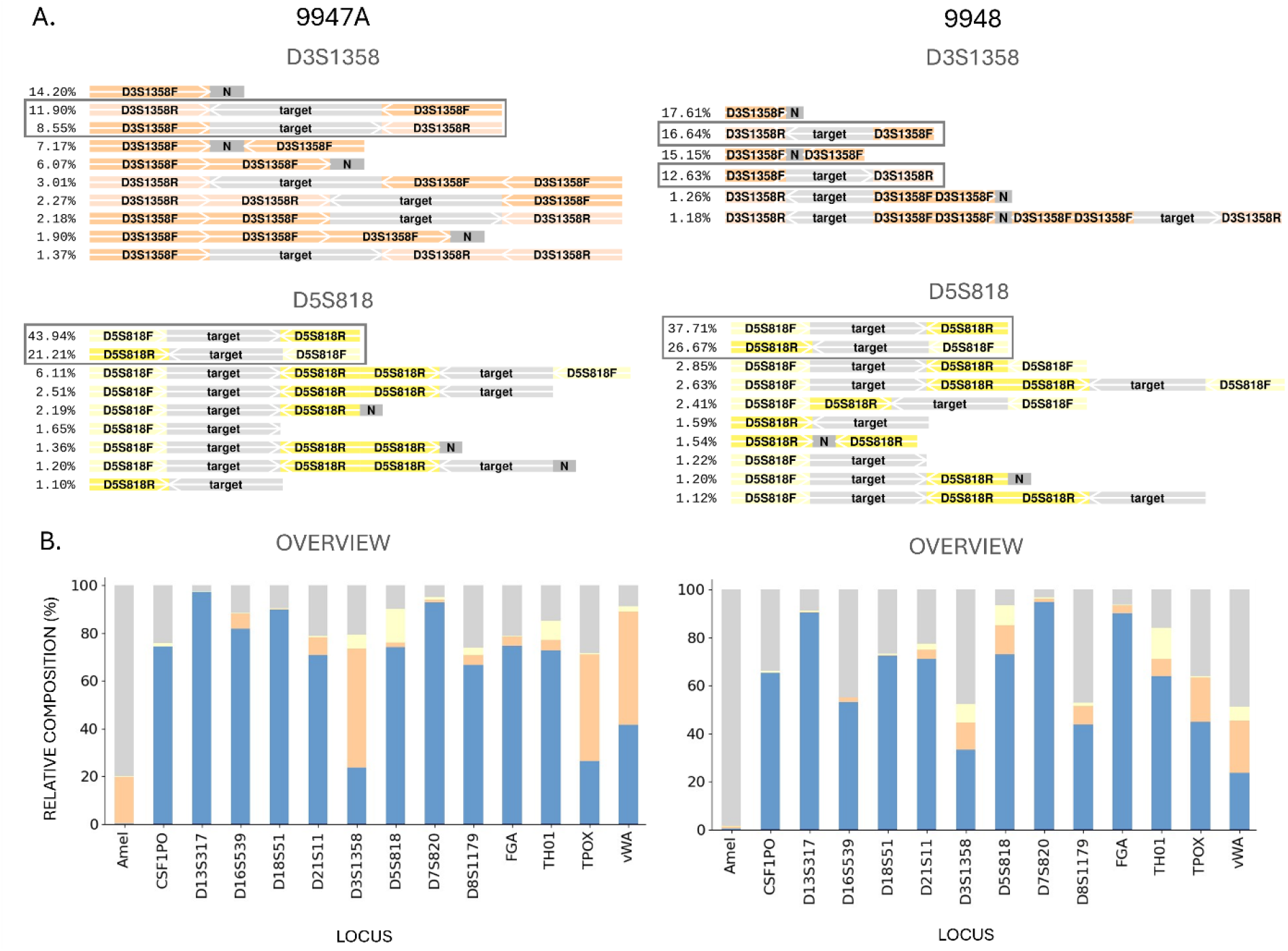
**A)** Readsaber summary figures of two representative loci, D3S1358 and D5S818, as given for sample 9947A and 9948. Forward and reverse primers are indicated by locusname + F or R, respectively. The target is indicated in grey and labelled as ‘target’. Grey blocks labelled as ‘N’ are sequences of 10 bp or longer without annotation. The relative read count per annotation pattern (%) is given at the left side of the annotation patterns. The arrows indicate the orientation of the primer and target sequences. Patterns representing true amplicons are framed. **B)** Overview of the target and artefact proportions of 14 loci, after amplification of 9947A and 9948.

The Readsaber plots revealed three additional structural features. First, native barcodes frequently remained present after trimming, with the highest rates seen for NB19 (Supplementary Figure S3-S4). This pattern may result from double barcode ligation during library preparation followed by incomplete barcode trimming as performed by Dorado. This phenomenon was recently described by Beeloo et al. (2025), who reported residual barcode sequences in approximately 7% of reads, with a barcode dependent variation in prevalence (28). As residual barcodes did not interfere with the identification and classification of RPA artefacts in the present study, no further trimming optimization was performed. Second, we often observed reads where either the first primer at the 5’ end, or the final primer at the 3’ end was not annotated, resulting in partial amplicons or amplicon multimers, as illustrated for locus D5S818 in Figure 2A. Detailed inspection of representative reads showed that these reads often contained a partial primer fragment. Although premature RPA termination due to polymerase dissociation cannot be excluded, these products were more likely attributable to low sequencing quality at the end of the reads, or incorrect trimming by Dorado as described above. Therefore these partial amplicons and partial amplicon multimers were grouped together with complete amplicons and amplicon multimers. Finally, sequences annotated as “N” were observed for almost all loci (Figure 2A). These elements were defined as stretches of at least 10 bp that did not match any predefined structural element in the annotation file or STR reference library. This feature was particularly pronounced for Amelogenin, resulting in low target proportions of 0.24% for 9947A and 0.34% for 9948 (Supplementary Figure S1). To investigate the origin of these unannotated sequences, representative reads were queried through NCBI blastn (29). The results indicated that these reads were mainly attributable to nonspecific amplification, with the strongest alignments to chromosome 6 of the homo sapiens reference genome. As primer design for Amelogenin is constrained by known single nucleotide polymorphisms (SNP) in the AMELX and AMELY regions, no further primer optimization was performed for this locus. Moreover, our previous study showed accurate genotyping for the same primers when combined with target-specific exo probes (6).

### 3.2. Comparison to PCR amplification

To verify whether these multimeric artefacts originate during RPA amplification, rather than during ONT library preparation, sequencing, or downstream data analysis, Readsaber was applied to sequencing data published by Tytgat et al. in 2022 (30), where PCR was used to amplify the same 14 forensic loci. It should be noted that a direct comparison between RPA and PCR in terms of accuracy, efficiency, or specificity is not appropriate, as RPA primers differ from PCR primers and are typically longer (36 bp). Nevertheless, the PCR dataset by Tytgat et al. provides a useful reference to evaluate whether comparable multimeric structures are also observed after amplification of the same loci with a non-RPA method. Supplementary Figure S5 shows the Readsaber summary figures of 9947A after PCR amplification. Across 14 analyzed loci, amplicon multimers accounted on average for 0.14% of reads (SD = 0.23), and primer multimers for 0.04% of reads (SD = 0.11), showing substantially lower multimer levels compared to our RPA data for the same sample, further supporting the conclusion that the multimers observed in this study are primarily RPA derived.

### 3.3. Temperature optimization

The effects of key reaction parameters on multimer formation was evaluated for sample 3 by Fragment Analysis. Two representative loci were amplified while varying one RPA parameter at a time: D3S1358 for primer multimers and D5S818 for amplicon multimers. Reaction temperature had the strongest effect on amplification specificity and efficiency. As shown in Figure 3A, lower temperatures generally reduced the fractions of both primer and amplicon multimers, while the absolute target concentration remained stable or even slightly increased. Figure 3B compares the Fragment Analyzer profiles after amplification at 42°C and 34°C. At 42°C, both loci showed multiple artefact peaks of which the size distribution corresponded to the multimeric products observed with ONT sequencing. In contrast, amplification at 34°C drastically improved specificity, increasing the target mass fraction from 15% to 83% for D3S1358 and from 84% to 98% for D5S818 (Figure 3A).

**Figure 3:**
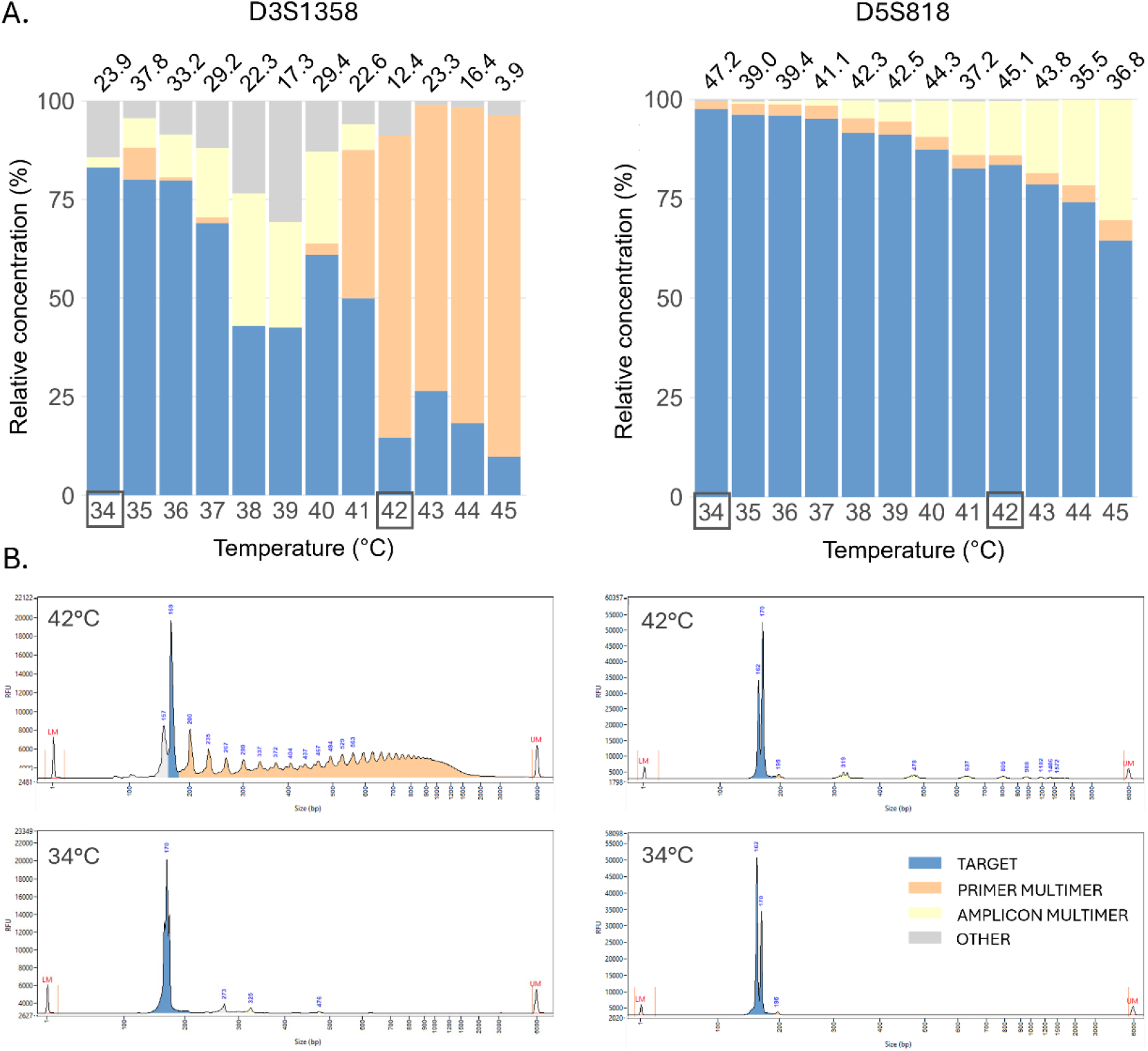
Temperature assessment for locus D3S1358 (true genotype 16,16) and D5S818 (true genotype 11,13). **A)** Relative target and multimer proportions across different temperatures. Bars represent the relative composition of each reaction, and the absolute target concentration (ng/µL) is given above each bar. **B)** Electropherograms obtained from Fragment Analyzer after RPA amplification at 42°C and 34°C. The electropherograms correspond to the reactions at 42°C and 34°C in panel A. Target proportions are indicated in blue, while primer multimer and amplicon multimer proportions are shown in orange and yellow, respectively. All non-identified RPA artefacts classified as ‘other’ are indicated in grey.

To determine whether these findings could be extrapolated to other loci, sample 9947A and 9948 were amplified for 14 loci at 34°C, and subsequently analyzed by ONT sequencing. Figure 4A compares the relative target and multimer read fractions after amplification at 42°C and 34°C. The Readsaber summary graphs are shown in Supplementary Figures S1 and S4, respectively. For each locus, the relative target and multimer read proportions at 42°C were subtracted from the corresponding proportions at 34°C. Overall, lowering the reaction temperature increased the relative target proportion for multiple loci. Among the loci showing an increase, the mean increase was 24.28 percentage points for 7/14 loci in sample 9947A and 25.22 percentage points for 9/14 loci in sample 9948. The relative target proportion was classified as stable, defined as an absolute change of no more than 5 percentage points, for 5/14 loci in 9947A and 2/14 loci in 9948. The relative target proportion decreased at 34°C for CSF1PO and TPOX in both samples, and for FGA in sample 9948. In all cases except TPOX in sample 9947A, this was attributable to increased off-target amplification rather than increased multimer formation. The largest improvements were observed for loci with substantial multimer formation at 42 °C, including D3S1358, D5S818, D8S1179, TH01, and vWA, which showed an average increase in relative target proportion of 40.81, 21.83, 35.20, 28.21, and 31.90 percentage points, respectively. Notably, reducing multimer formation at 34°C did not substantially increase genotyping accuracy (Figure 4B). At both temperatures, single allelic dropouts were observed at four loci, while one additional locus showed a double allelic dropout at 42°C. Most dropouts occurred for Amelogenin, likely because nonspecific amplification reduced the number of target reads available for reliable genotyping. As sequencing allows individual reads to be classified and excluded according to their sequence, multimer reduction may have a greater impact on genotyping accuracy for other detection methods, such as electrophoresis or intercalating dye-based assays, where multimers can overlap with target products or contribute directly to the fluorescent signal.

**Figure 4:**
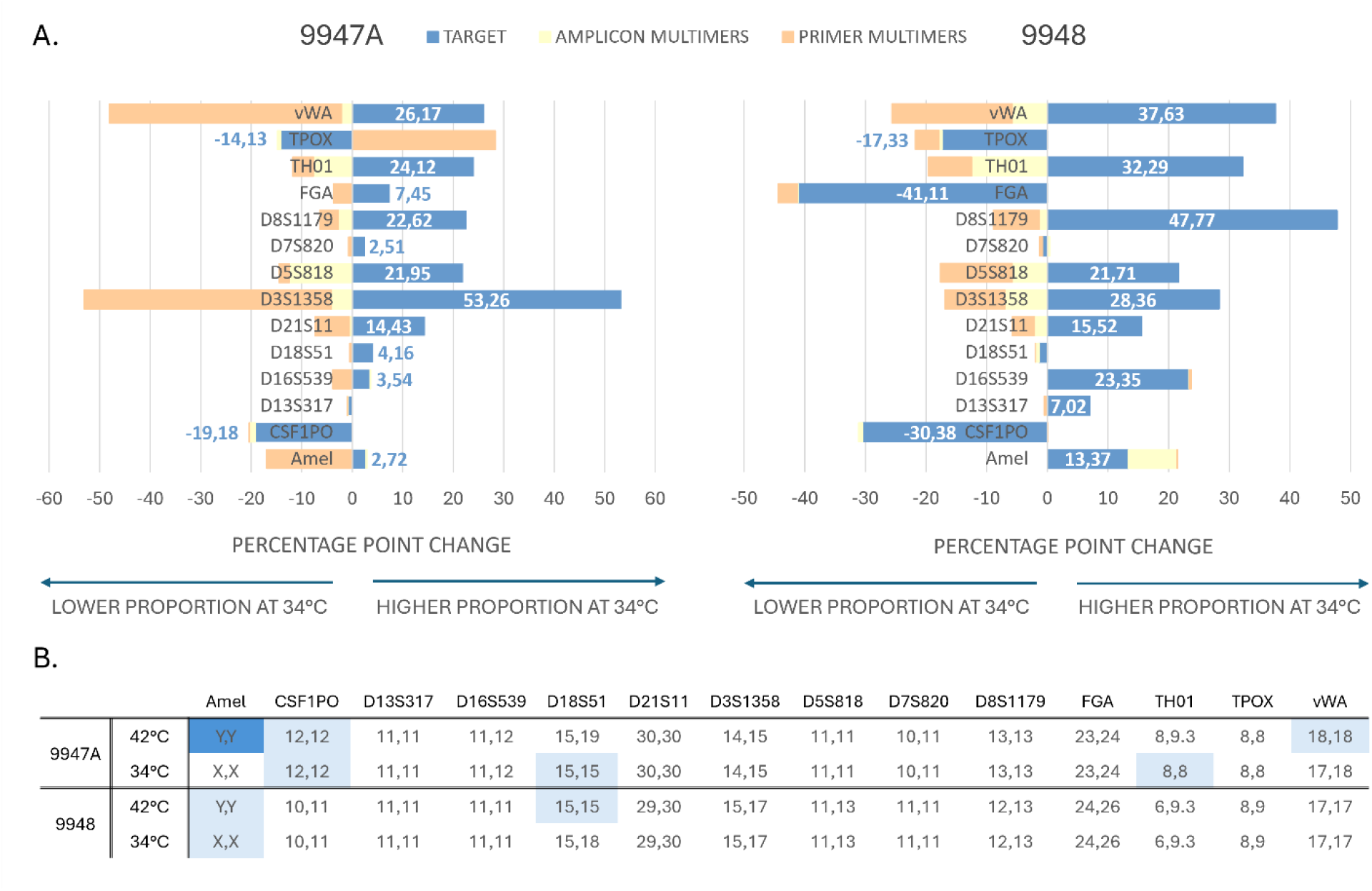
**A)** Change in relative target and multimer read proportions after amplification at 34°C compared to 42°C. Changes are expressed in percentage points and were calculated as the relative read proportion at 34°C minus the corresponding proportion at 42°C. Positive values indicate an increased relative proportion at 34°C, whereas negative values indicate a decreased relative proportion at 34°C. **B)** Genotyping accuracy at both temperatures. Correctly genotyped loci are marked in white. Loci with one allelic dropout are indicated in light blue, while loci with two incorrect alleles are indicated in dark blue.

Although the relative target proportion either remained stable or increased for most loci at 34°C, overall reaction efficiency was reduced, as the total post-amplification concentration (comprising both target and artefact products) was frequently lower (Supplementary Table S3). This was particularly pronounced for D16S539 and D7S820. Both loci were characterized by low amplification efficiency at 42°C, with average post-amplification total concentrations of 5.73 ng/µL for D16S539 and 5.09 ng/µL for D7S820. At 34°C these concentrations further decreased to 0.13 ng/µL for D16S539 and 0.17 ng/µL for D7S820. Importantly, a reduction in total concentration does not necessarily correspond to a reduction in absolute target concentration, as described above. No association was observed between reaction efficiency or specificity and amplicon length or GC content (Supplementary Table S2). Overall, these findings indicate that lowering the amplification temperature might improve reaction specificity particularly for loci prone to multimer formation. However, this strategy appears less suitable for loci with low amplification efficiency or substantial nonspecific amplification, as it may further reduce total product yield while maintaining off-target amplification.

### 3.4. Optimization of other reaction conditions

Beyond reaction temperature, primer concentration and T4 UvsX concentration were the most influential parameters. The Fragment Analyzer profiles showed that lower primer concentrations and higher T4 UvsX concentrations reduced the relative proportion of both primer and amplicon multimers (Figure 5). Notably, doubling the T4 UvsX concentration from 0.06 to 0.12 mg/mL increased the absolute target concentration twofold for D3S1358 and 1.3-fold for D5S818, highlighting its importance for amplification efficiency. In contrast, reducing the primer concentration showed an inverse relationship between relative and absolute target concentration, indicating a trade-off between reaction specificity and efficiency for this parameter. Moreover, lower primer concentrations ranging from 0.1-0.3 µM led to allelic dropouts for D5S818. Reducing the Bst polymerase concentration lead to a small increase in relative target proportion, although an allelic drop-in occurred at D3S1358 when polymerase concentration was reduced to 0.015 U/µL. No substantial differences in multimer formation or target yield were observed across the tested dNTP, Mg^2+^, T4 UvsY, and GP32 concentrations. Increasing the DNA input substantially reduced primer multimer proportions, whereas amplicon multimer formation remained largely stable across the tested input range. This is probably caused by the increased probability of primer-template interactions at higher DNA inputs, thereby reducing primer-primer interactions. In contrast, amplicon multimer formation may be less affected by initial template concentration, as the final amplicon yield and thus the final amplicon multimer yield is limited by reagent availability during the RPA reaction. For both loci, the Fragment Analyzer profiles of the no template controls (NTC) consisted exclusively of peaks with size spacing consistent with primer multimers (Supplementary Figure S6). While the presence of primer multimers was not confirmed by sequencing, the fragments suggest that their formation is a template independent process. Despite the absence of target DNA in both NTCs, the total post-amplification DNA concentrations reached 63.10 ng/µL for D3S1358 and 52.52 ng/µL for D5S818. These findings indicate that primer multimer formation may lead to false-positive results, for instance when nonspecific fluorescent dyes are used for detection.

**Figure 5:**
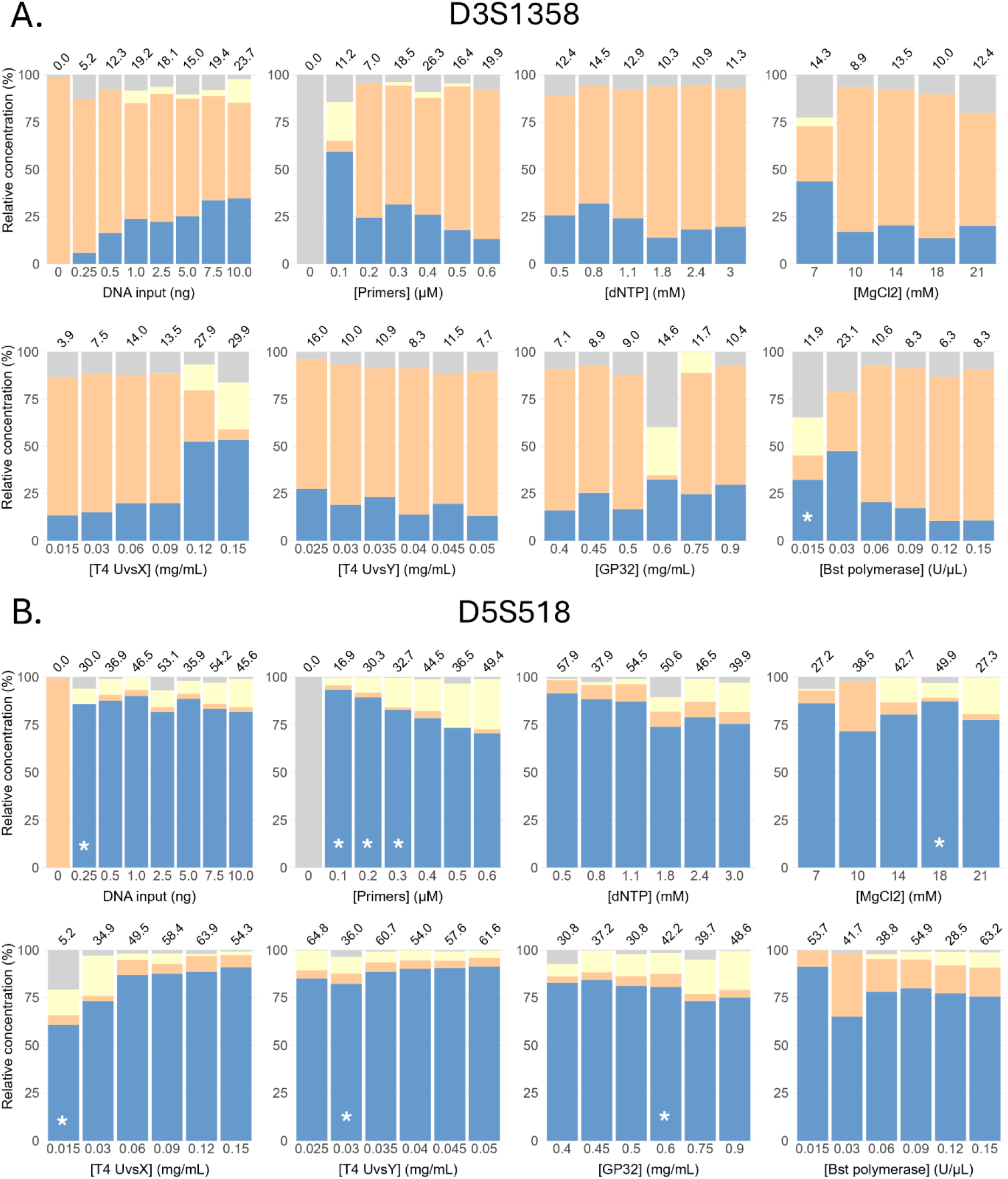
Overview of the target, primer and amplicon multimer fractions for **A)** locus D3S1358 (true genotype 16,16) and **B)** locus D5S818 (true genotype 11,13) under variable reaction conditions. The bars show the relative composition for each condition, and the absolute target concentration (ng/µL) is given above each bar. Allelic dropouts are indicated by a star.

## 4. Discussion

For the past two decades, RPA has gained increasing interest as an isothermal alternative to PCR. The ability to amplify nucleic acids at low and constant temperatures makes RPA attractive for applications in low-resource environments and portable diagnostic platforms (5). However, nonspecific artefact formation, including off-target amplification, ab initio DNA synthesis, and primer dimer or hairpin formation, remains an important limitation of this technique (13). This study identified two new types of RPA artefacts, which we refer to as primer multimers and amplicon multimers. Unlike conventional primer dimers, which are generally short and easily distinguished from target amplicons by electrophoresis, multimeric artefacts typically generate longer products that may overlap in length with target amplicons. Besides, once formed, both primer multimers and amplicon multimers contain multiple primer hybridization sites, which further result in the exponential amplification of heterogeneous byproducts and smear formation. This is particularly relevant in multiplex assays, where multimers derived from one locus, may fall within the expected size range of the target amplicons from another locus.

RPA byproducts may also affect real-time detection with intercalating dyes, especially in the absence of target DNA. For instance, our NTC results showed high-molecular-weight artefacts consistent with primer multimers, which contributed to high total DNA concentrations. These products may lead to false-positive results due to nonspecific binding of the dye. Although not evaluated in this study, we reason that target-specific probes, such as exo probes, may help mitigate this issue. Exo probes generate fluorescence only after hybridization to their complement (31). Therefore, primer multimers lacking the corresponding target region would not be expected to generate a fluorescent signal. This hypothesis is supported by our previous study (6), in which the same Amelogenin primers were combined with target-specific exo probes for sex typing, without false-positive signals in any of the NTCs. In contrast, such probes are unlikely to distinguish true amplicons from amplicon multimers, as both contain target sequences. Since amplicon multimers are target-derived products, their presence may not compromise qualitative diagnostic readouts, provided that the assay only aims to detect the presence or absence of a target sequence. However, as the kinetics of amplicon multimer formation have not yet been fully characterized, these products could potentially affect quantitative RPA applications. Although quantitative RPA assays have been successfully established (15,32–34), these assays generally distinguish target concentrations over ten- to hundredfold ranges. This indicates that further studies are necessary to establish to what extent amplicon multimers might influence assay precision. This concern is consistent with findings by Garafutdinov et al. (2020), who experienced scattered time-to-threshold (T_t_) values between replicates when investigating Bst polymerase induced DNA multimerization. They attributed this variability to the stochastic and rare initiation of multimerization, where the onset of multimer formation can vary substantially even between replicate reactions (35).

To the best of our knowledge, such multimer formation has not yet been systematically described for RPA reactions. However, similar products have been reported in other isothermal amplification assays driven by strand displacing polymerases. For example, Hafner et al. (2001) described linear target isothermal multimerization amplification (LIMA), an isothermal method using two target derived primers that generated ladder-like electrophoretic bands corresponding to products larger than the expected amplicon (36). Wang et al. (2017) later proposed that LIMA is part of a broader phenomenon called unusual isothermal multimerization and amplification (UIMA), which can occur with only one primer (37). Both mechanisms have been proposed to occur as a byproduct in other isothermal assays including loop-mediated isothermal amplification (LAMP), and are assumed to involve the formation of cyclic DNA intermediates followed by extension of self-annealed 3′ ends. However, the precise molecular mechanism remains unresolved (37,38). For target-derived DNA multimerization, the presence of both double-stranded DNA and at least one primer has been proposed to be required. In contrast, Bst polymerase has also been reported to generate nonspecific amplification products in the absence of target and primers, for example through ab initio DNA synthesis (35). Such template independent activity could potentially contribute to the primer multimers observed in the NTC samples in the present study. The isothermal assays described above rely on strand displacing DNA polymerases, often Bst polymerase, which is also used in Invitrogen’s RPA kit in the present study. In our experiments, lower Bst polymerase concentrations appeared to reduce multimer formation to some extent. However, further studies including the systematic variation of polymerase type and activity, would be required to assess to which extent these factors contribute to the artefacts observed here.

Overall, reaction temperature, primer concentration and T4 UvsX concentration had the strongest effects on multimer formation. Of these parameters, only reaction temperature was investigated in greater detail with ONT sequencing, as reducing the primer concentration decreased overall yield and caused allelic dropouts. Although increasing T4 UvsX concentration may offer an interesting optimization strategy, this approach comes with a substantial increase in reagent cost. The temperature assessment showed that the relative target amplicon concentration reached a maximum at an amplification temperature of 34°C. The temperature range evaluated in this study was selected based on Invitrogen’s recommended range of 34– 45°C. In theory, however, the optimal target proportion may occur at even lower temperatures. It should also be noted that amplification temperature may interact with reaction time, which was kept at 40 minutes across all temperatures. At 34°C, lower reaction kinetics may have delayed the formation of RPA artefacts, resulting in a higher relative target concentration compared with higher temperatures. An alternative strategy could be to reduce the reaction time at higher temperatures, which may potentially limit artefact formation in a similar way as reducing reaction temperature.

The concentration of single-stranded DNA-binding protein (GP32) only had a minor effect on absolute target yield and did not substantially affect the relative target concentration. Interestingly, this observation differs from the findings of Cordoba-Andrade et al. (2025), who reported a twofold increase in RPA product yield for a four-fold increase in GP32 concentration (16). This discrepancy might be explained by differences in assay design, target sequence, or reaction composition. For instance, the effect of GP32 may depend on interactions with other RPA reagents, which have not been investigated in this study. In addition, the difference in artefacts may also have contributed to the observed differences, as Cordoba-Andrade et al. focused on primer dimer formation, whereas the present study investigates multimeric artefacts. Nevertheless, the reported influence of the RPA enzymes on reaction specificity and efficiency emphasizes the need for greater transparency regarding the composition of RPA reagents. Although Twistdx (Maidenhead, UK) remains one of the largest suppliers of RPA reagents, its kits rely on pre-assembled reagent mixes with undisclosed enzyme and buffer compositions. In contrast, newer RPA formulations, such as the RPA enzymes from Watchmaker Genomics (Boulder, CO, USA) and the Lyo-ready RPA kit by Invitrogen (Waltham, MA, USA) used in this study, fully disclose reagent compositions and concentrations, allowing for systematic optimization of individual components.

Given the limited number of loci included in this proof-of-concept study, no definitive conclusions could be drawn regarding which primer or target features promote or prevent multimer formation. However, inspection of the junctions revealed a recurrent structural pattern among primer multimers. Across all loci showing primer multimer formation, primers were predominantly arranged as head-to-tail repeats, with deletions of 1–3 bp frequently observed at the repeat junctions. This pattern is consistent with findings by Garafutdinov et al. (2020), who reported multimers for isothermal amplification with Bst exo− DNA polymerase, consisting of junctional deletions of 2–4 nucleotides (35). For amplicon multimers, the repeat junctions were typically consistent with a blunt-end 5′–5′ junction between identical primers. To the best of our knowledge, such a multimeric pattern consisting of multiple amplicons has not been described in literature before.

In this study, we aimed to identify, characterize, and mitigate the RPA artefacts arising during the amplification of forensic STR loci. Improving reaction specificity is essential for forensic genetics, where DNA input amounts are generally low and nonspecific products may interfere with accurate genotyping and downstream profile interpretation. More broadly, this study is, to the best of our knowledge, one of the first to systematically assess the influence of RPA enzyme concentrations on amplification efficiency and specificity. Although presented as a proof-of-concept study, our findings highlight the need for a more fundamental understanding of the RPA reaction and its potential side products. Such knowledge is relevant beyond forensic genetics, for example in rapid diagnostics, genetic screening, biosensing, and other fields where RPA is used as a rapid isothermal amplification method. Future work should investigate how the RPA mechanism and its components affect key assay parameters, including specificity, sensitivity, efficiency, robustness, and reproducibility. By opening up the discussion for more fundamental research into the RPA mechanism, this paper may support such studies towards the development of more reliable and interpretable RPA workflows.

## 5. Conclusion

This study identified and characterized two previously undescribed RPA-derived artefacts, referred to as primer multimers and amplicon multimers, during the amplification of forensic STR loci. ONT sequencing and fragment analysis showed that these products consist of repeated primer or amplicon units and originate during RPA amplification rather than during sequencing or downstream analysis. Among the tested reaction conditions, temperature had the strongest effect on artefact formation, with amplification at 34 °C reducing multimer formation compared to 42 °C. Primer concentration and T4 UvsX concentration also influenced the balance between target yield and reaction specificity. Although the optimized conditions reduced artefact formation for multiple loci, the effect remained locus-dependent. Overall, this proof-of-concept study provides new insight into RPA artefact formation and highlights the importance of systematic reaction optimization. While demonstrated in a forensic STR context, these findings may support the development of more reliable RPA workflows for broader applications, including rapid diagnostics, biosensing, and portable molecular testing.

## Acknowledgements

We thank Peter De Rijk from the Center for Molecular Neurology at the University of Antwerp for his valuable assistance and expertise in the use of Readsaber.

We also thank David Van Hoofstat from Eurofins Forensics Belgium NV for performing the sample preparation and data analysis for benchmark genotyping of the samples.

Finally, we thank Sarah De Keulenaer from the Ghent University sequencing core facility NXTGNT for her assistance with Oxford Nanopore Technologies sequencing.

## Data availability statement

The sequencing datasets used in this study may be found at NCBI-SRA. The names and accession numbers are listed at https://www.ncbi.nlm.nih.gov/bioproject/PRJNA1505172.

## Author contributions

LDK: Conceptualization, Methodology, Validation, Formal analysis, Investigation, Data curation, Writing – original draft, Visualization, Project administration. KD: Methodology, Formal analysis, Data curation, Writing – review and editing. SS: Conceptualization, Methodology, Writing – review and editing. DVH: Formal analysis, Investigation, Writing – review and editing. DD: Methodology, supervision, Writing – review and editing. FVN: Conceptualization, Methodology, Supervision, Writing – review and editing.

