## Supplementary materials for "Recombinase polymerase amplification: characterization and mitigation of undescribed multimeric artefacts"

### 1. TABLES

**Table S1:** Overview of the samples and their true genotypes.

| | Amel | CSF1PO | D13S317 | D16S539 | D18S51 | D21S11 | D3S1358 | D5S818 | D7S820 | D8S1179 | FGA | TH01 | TPOX | vWA |
|---|---|---|---|---|---|---|---|---|---|---|---|---|---|---|
| S1 (9947A) | X,X | 10,12 | 11,11 | 11,12 | 15,19 | 30,30 | 14,15 | 11,11 | 10,11 | 13,13 | 23,24 | 8,9.3 | 8,8 | 17,18 |
| S2 (9948) | X,Y | 10,11 | 11,11 | 11,11 | 15,18 | 29,30 | 15,17 | 11,13 | 11,11 | 12,13 | 24,26 | 6,9.3 | 8,9 | 17,17 |
| S3 | X,X | 7,11 | 12,12 | 11,12 | 11,16 | 30,30.2 | 16,16 | 11,13 | 9,10 | 10,14 | 19,24 | 6,9.3 | 9,11 | 16,17 |

**Table S2:** Overview of the RPA primers and amplicon properties.
The amplicon length and GC content are given as the range between the shortest and longest allele in the reference library containing all known STR alleles present in more than 1% of the world population, as collected from NIST STRbase.

| LOCUS | PRIMER | SEQUENCE | PRIMER LENGTH (NT) | AMPLICON LENGTH (BP) | AMPLICON GC CONTENT (%) |
|---|---|---|---|---|---|
| Amelogenin | F | CCCTGGGCTCTGTAAAGAATAGTG | 24 | 112-118 | 44.9-44.6 |
| | R | CCAACCATCAGAGCTTAAACTGGGAAGCTG | 30 | | |
| CSF1PO | F | TTGGACAGCATTTCCTGTGTCAGACCCTGTTC | 32 | 179-199 | 37.4-36.2 |
| | R | GGACTAGCAGGTTGCTAACCACCCTGTGTCTC | 32 | | |
| D3S1358 | F | CATCTCTTATACTCATGAAATCAACAGAGGCTTGC | 35 | 149-173 | 40.9-39.3 |
| | R | GAGCTATGATTCCCCCACTGCAGTCCAATCTGGGT | 35 | | |
| D5S818 | F | TATGTGACAAGGGTGATTTTCCTCTTTGGTATCC | 34 | 147-171 | 29.9-29.2 |
| | R | TCCAATCATAGCCACAGTTTACAACATTTGTATCT | 35 | | |
| D7S820 | F | ATAACGATTCCACATTTATCCTCATTGAC | 29 | 233-269 | 30.0-29.4 |
| | R | GGTTTCACCATGTTGGTCAGGCTGACTATG | 30 | | |

| LOCUS | PRIMER | SEQUENCE | PRIMER LENGTH (NT) | AMPLICON LENGTH (BP) | AMPLICON GC CONTENT (%) |
|---|---|---|---|---|---|
| D8S1179 | F | CTGGCAACTTATATGTATTTTTGTATTTCATG | 32 | 214-246 | 30.8-30.5 |
| | R | TTTTACCAAATTGTGTTCATGAGTATAGTTTC | 32 | | |
| D13S317 | F | TGGTATCACAGAAGTCTGGGATGTGGAGGA | 30 | 195-219 | 40.0-38.4 |
| | R | GTTGAGCCATAGGCAGCCCAAAAAGACAGA | 30 | | |
| D16S539 | F | GGGGGTCTAAGAGCTTGTAAAAAG | 24 | 276-300 | 39.9-38.7 |
| | R | GTTTGTGTGTGCATCTGTAAGCATGTATC | 29 | | |
| D18S51 | F | GGCAGGAGGAGTTCTTGAGCCCAGAAGGTTA | 31 | 312-360 | 39.7-37.8 |
| | R | ACCCGACTACCAGCAACAACACAAATAAAC | 30 | | |
| D21S11 | F | CTCCATAAATATGTGAGTCAATTCCCCAAG | 30 | 227-263 | 33.9-32.7 |
| | R | ATGTTGTATTAGTCAATGTTCTCCAGAGAC | 30 | | |
| FGA | F | ATTCATGGAAGGCTGCAGGGCATAACATTA | 30 | 345-385 | 35.7-34.8 |
| | R | TACTTTTTCTATGACTTTGCGCTTCAGGA | 29 | | |
| TH01 | F | TATCTGGGCTCTGGGGTGATTCCCATTGGCCTGTTC | 36 | 191-207 | 53.9-51.7 |
| | R | GCACCGAAGACCCCTCCTGTGGGCTGAAAAGCTC | 34 | | |
| TPOX | F | CCAGAACCGTCGACTGGCACAGAACAGGCACTTAGG | 36 | 292-308 | 50.3-49.0 |
| | R | TCGTGTTTGCGTCCCCAACGCTCAAACGTGAGGTTG | 36 | | |
| vWA | F | GCCCTAGTGGATGATAAGAATAATCAGTATGTG | 33 | 139-167 | 36.0-34.1 |
| | R | GGACAGATGATAAATACATAGGATGGATGG | 30 | | |

**Table S3:** Comparison of total read count and total post-amplification DNA concentration after amplification at 42°C and 34°C. Total concentrations are measured by Qubit (ng/µL). Total values include both target and artefact read counts or concentrations. The effect of temperature is expressed as the ratio between the values obtained at 34°C and 42°C.

## 9947A

| | CONCENTRATION (ng/µL) | | | READ NUMBER | | |
|---|---|---|---|---|---|---|
| | 42°C | 34°C | Ratio (34/42) | 42°C | 34°C | Ratio (34/42) |
| Amelogenin | 8.84 | 5.62 | 0.64 | 334 457 | 203 313 | 0.61 |
| CSF1PO | 24.30 | 6.04 | 0.25 | 178 999 | 130 695 | 0.73 |
| D3S1358 | 22.10 | 20.40 | 0.92 | 327 999 | 365 211 | 1.11 |
| D5S818 | 29.90 | 19.90 | 0.67 | 146 598 | 228 829 | 1.56 |
| D7S820 | 6.32 | 0.11 | 0.02 | 720 344 | 27 400 | 0.04 |
| D8S1179 | 35.20 | 1.63 | 0.05 | 194 737 | 161 327 | 0.83 |
| D13S317 | 35.30 | 14.50 | 0.41 | 319 784 | 190 714 | 0.60 |
| D16S539 | 6.18 | 0.10 | 0.02 | 228 494 | 8 091 | 0.04 |
| D18S51 | 35.20 | 12.40 | 0.35 | 182 948 | 99 327 | 0.54 |
| D21S11 | 42.90 | 17.00 | 0.40 | 853 726 | 299 534 | 0.35 |
| FGA | 30.00 | 0.75 | 0.03 | 250 953 | 99 522 | 0.40 |
| TH01 | 25.40 | 13.60 | 0.54 | 134 537 | 126 445 | 0.94 |
| TPOX | 18.85 | 6.07 | 0.32 | 188 721 | 138 005 | 0.73 |
| vWA | 25.30 | 11.55 | 0.46 | 343 331 | 166 712 | 0.49 |

## 9948

| | CONCENTRATION (ng/µL) | | | READ NUMBER | | |
|---|---|---|---|---|---|---|
| | 42°C | 34°C | Ratio (34/42) | 42°C | 34°C | Ratio (34/42) |
| Amelogenin | 9.33 | 5.27 | 0.56 | 533 249 | 243 141 | 0.46 |
| CSF1PO | 18.50 | 6.84 | 0.37 | 281 563 | 166 088 | 0.59 |
| D3S1358 | 29.70 | 19.10 | 0.64 | 371 383 | 267 916 | 0.72 |
| D5S818 | 18.30 | 26.10 | 1.43 | 269 130 | 313 471 | 1.16 |
| D7S820 | 3.86 | 0.24 | 0.06 | 675 661 | 28 072 | 0.04 |
| D8S1179 | 30.00 | 1.48 | 0.05 | 223 250 | 171 587 | 0.77 |
| D13S317 | 33.90 | 10.40 | 0.31 | 450 692 | 221 355 | 0.49 |
| D16S539 | 5.27 | 0.15 | 0.03 | 250 993 | 9 111 | 0.04 |
| D18S51 | 32.00 | 7.14 | 0.22 | 245 348 | 120 013 | 0.49 |
| D21S11 | 21.10 | 15.25 | 0.72 | 1 006 488 | 641 908 | 0.64 |
| FGA | 17.00 | 1.26 | 0.07 | 369 101 | 165 847 | 0.45 |
| TH01 | 13.30 | 11.55 | 0.87 | 197 086 | 119 997 | 0.61 |
| TPOX | 10.60 | 4.12 | 0.39 | 385 339 | 176 069 | 0.46 |
| vWA | 15.25 | 7.48 | 0.49 | 220 330 | 162 800 | 0.74 |

### 2. Figures

**Figure S1:** Readsaber plots of sample 9947A and 9948 after RPA amplification at 42°C.
Forward and reverse primers are indicated by locusname + F or R, respectively. The target is indicated in grey and labelled as ‘target’. Grey blocks labelled as ‘N’ are sequences of 10 bp or longer without annotations. The relative read count per annotation pattern (%) is given at the left side of the annotation patterns. The arrows indicate the orientation of the primer and target sequences.

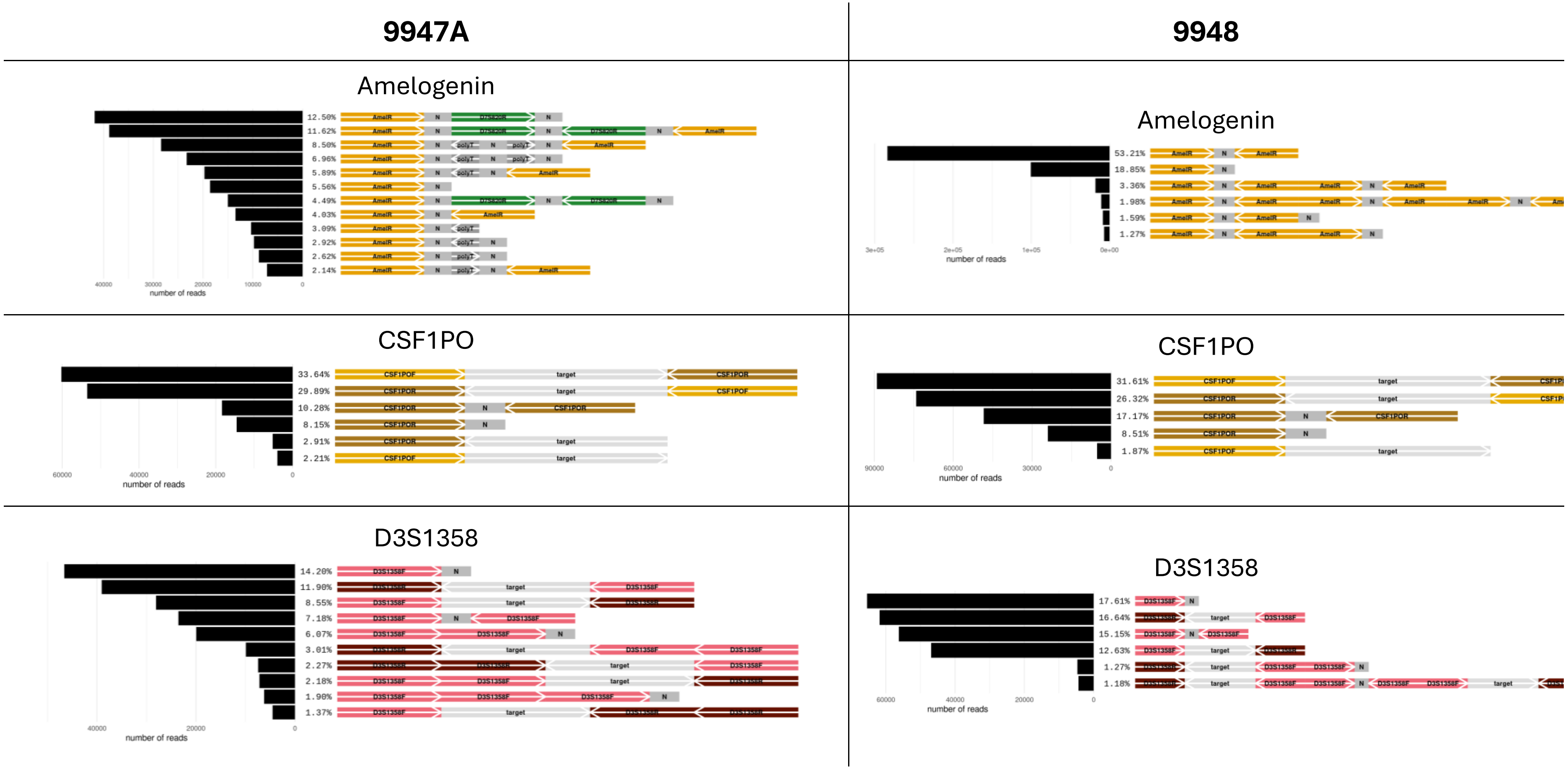

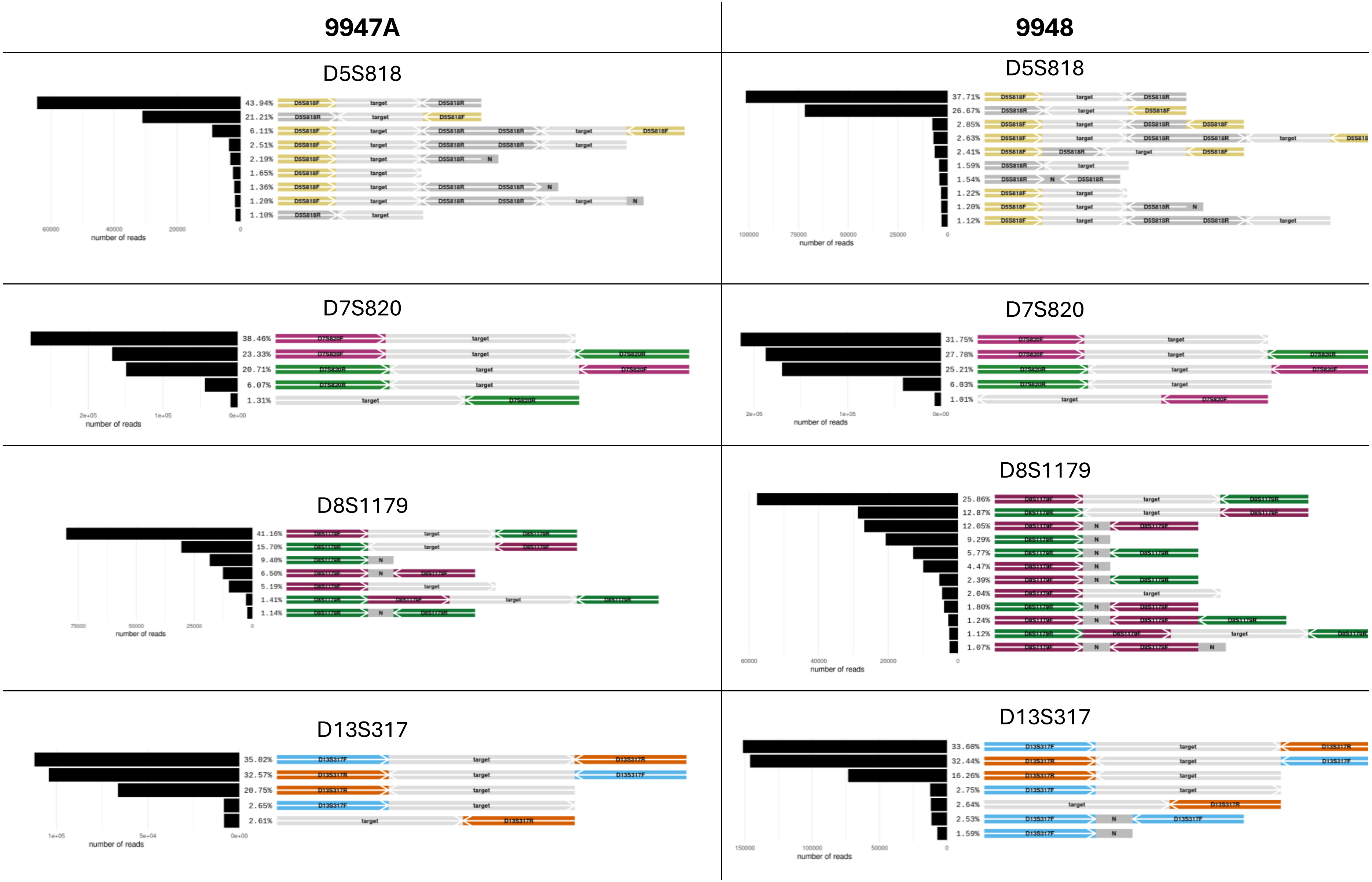
9947A
9948
D5S818
43.94%
21.21%
6.11%
2.51%
2.19%
1.65%
1.36%
1.20%
1.10%
number of reads
D5S818
37.71%
26.67%
2.85%
2.63%
2.41%
1.59%
1.54%
1.22%
1.20%
1.12%
number of reads
D7S820
38.46%
23.33%
20.71%
6.07%
1.31%
number of reads
D7S820
31.75%
27.78%
25.21%
6.03%
1.01%
number of reads
D8S1179
41.16%
15.70%
9.40%
6.50%
5.19%
1.41%
1.14%
number of reads
D8S1179
25.86%
12.87%
12.05%
9.29%
5.77%
4.47%
2.39%
2.04%
1.80%
1.24%
1.12%
1.07%
number of reads
D13S317
35.02%
32.57%
20.75%
2.65%
2.61%
number of reads
D13S317
33.60%
32.44%
16.26%
2.75%
2.64%
2.53%
1.59%
number of reads

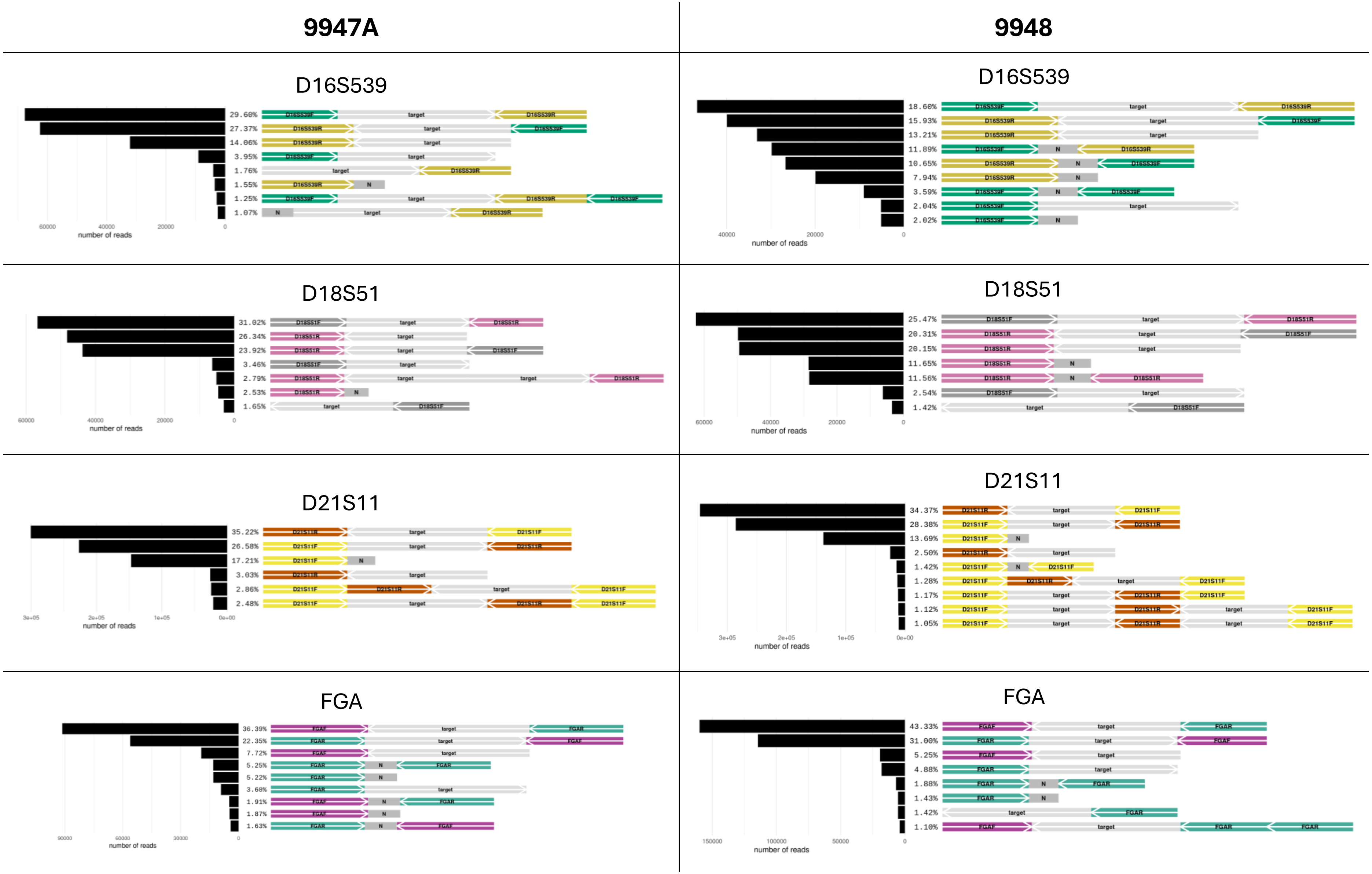
9947A
9948
D16S539
29.60%
27.37%
14.06%
3.95%
1.76%
1.55%
1.25%
1.07%
number of reads
D16S539
18.60%
15.93%
13.21%
11.89%
10.65%
7.94%
3.59%
2.04%
2.02%
number of reads
D18S51
31.02%
26.34%
23.92%
3.46%
2.79%
2.53%
1.65%
number of reads
D18S51
25.47%
20.31%
20.15%
11.65%
11.56%
2.54%
1.42%
number of reads
D21S11
35.22%
26.58%
17.21%
3.03%
2.86%
2.48%
number of reads
D21S11
34.37%
28.38%
13.69%
2.50%
1.42%
1.28%
1.17%
1.12%
1.05%
number of reads
FGA
36.39%
22.35%
7.72%
5.25%
5.22%
3.60%
1.91%
1.87%
1.63%
number of reads
FGA
43.33%
31.00%
5.25%
4.88%
1.88%
1.43%
1.42%
1.10%
number of reads

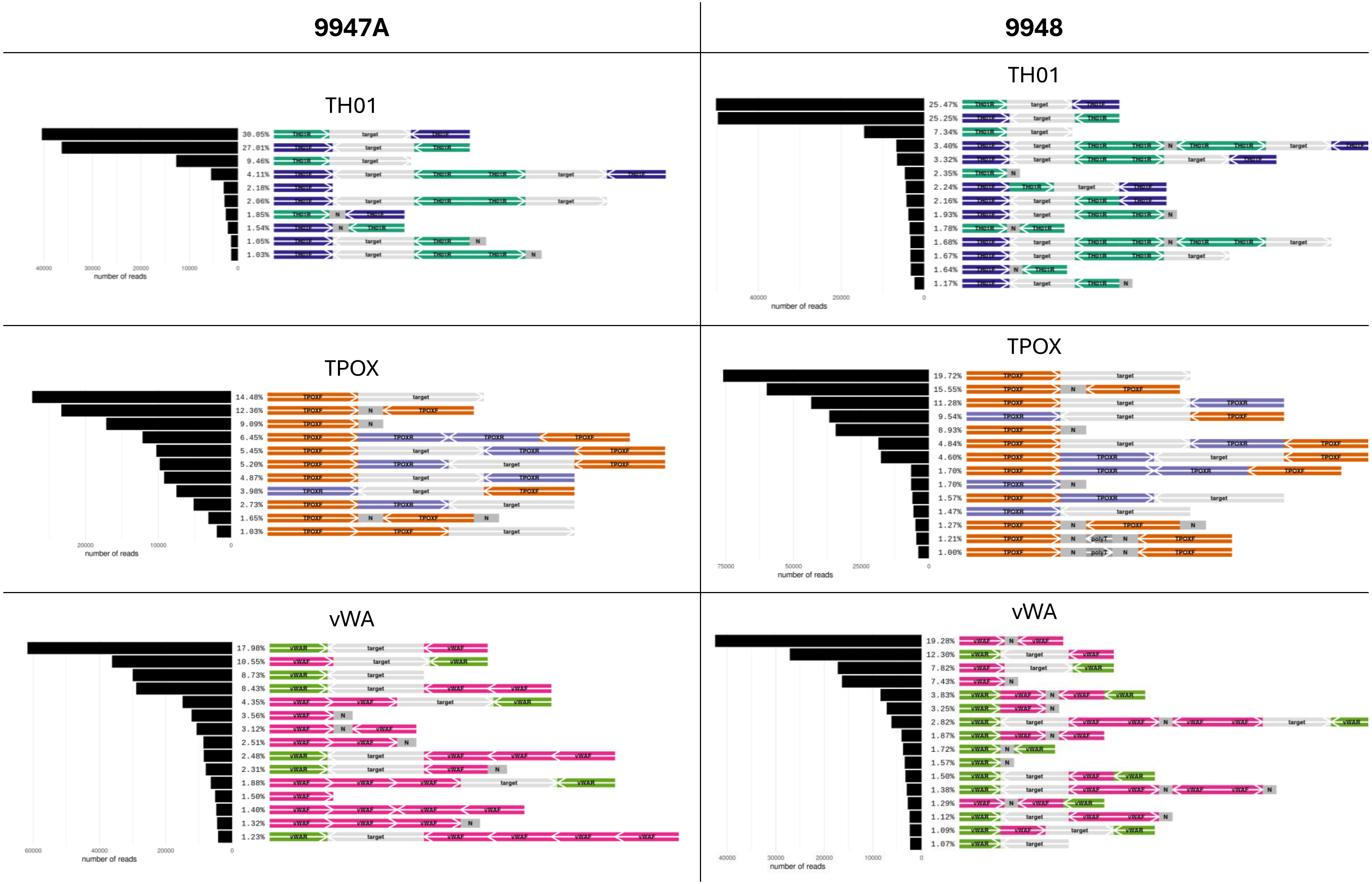
9947A
9948
TH01
30.05%
27.01%
9.46%
4.11%
2.18%
2.06%
1.85%
1.54%
1.05%
1.03%
number of reads
TH01
25.47%
25.25%
7.34%
3.40%
3.32%
2.35%
2.24%
2.16%
1.93%
1.78%
1.68%
1.67%
1.64%
1.17%
number of reads
TPOX
14.48%
12.36%
9.09%
6.45%
5.45%
5.20%
4.87%
3.98%
2.73%
1.65%
1.03%
number of reads
TPOX
19.72%
15.55%
11.28%
9.54%
8.93%
4.84%
4.60%
1.70%
1.70%
1.57%
1.47%
1.27%
1.21%
1.00%
number of reads
vWA
17.98%
10.55%
8.73%
8.43%
4.35%
3.56%
3.12%
2.51%
2.48%
2.31%
1.88%
1.50%
1.40%
1.32%
1.23%
number of reads
vWA
19.28%
12.30%
7.82%
7.43%
3.83%
3.25%
2.82%
1.87%
1.72%
1.57%
1.50%
1.38%
1.29%
1.12%
1.09%
1.07%
number of reads

**Figure S2:** Primer and target count per read for sample 9947A and 9948.

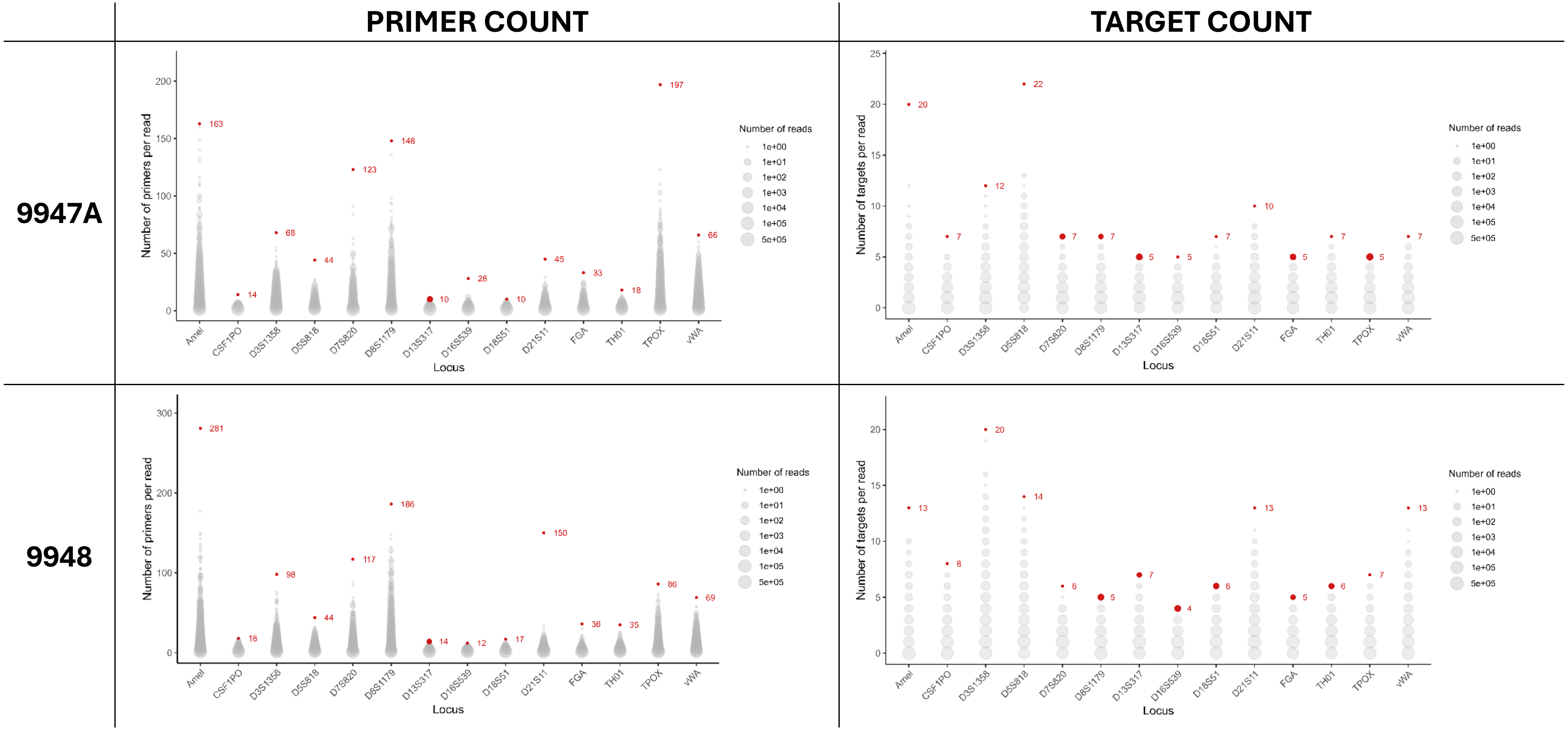

**Figure S3:** Overview of the remaining native barcodes (NB) in each sample. For each locus, the percentage represents the proportion of reads containing at least one native barcode. The y-axis is given from 0 to 5% to facilitate comparison at low proportions. NB17 and NB18 correspond to sample 9947A and 9948 following RPA amplification at 42°C, respectively. NB19 and NB20 were respectively used for sample 9947A and 9948 after amplification at 34°C.

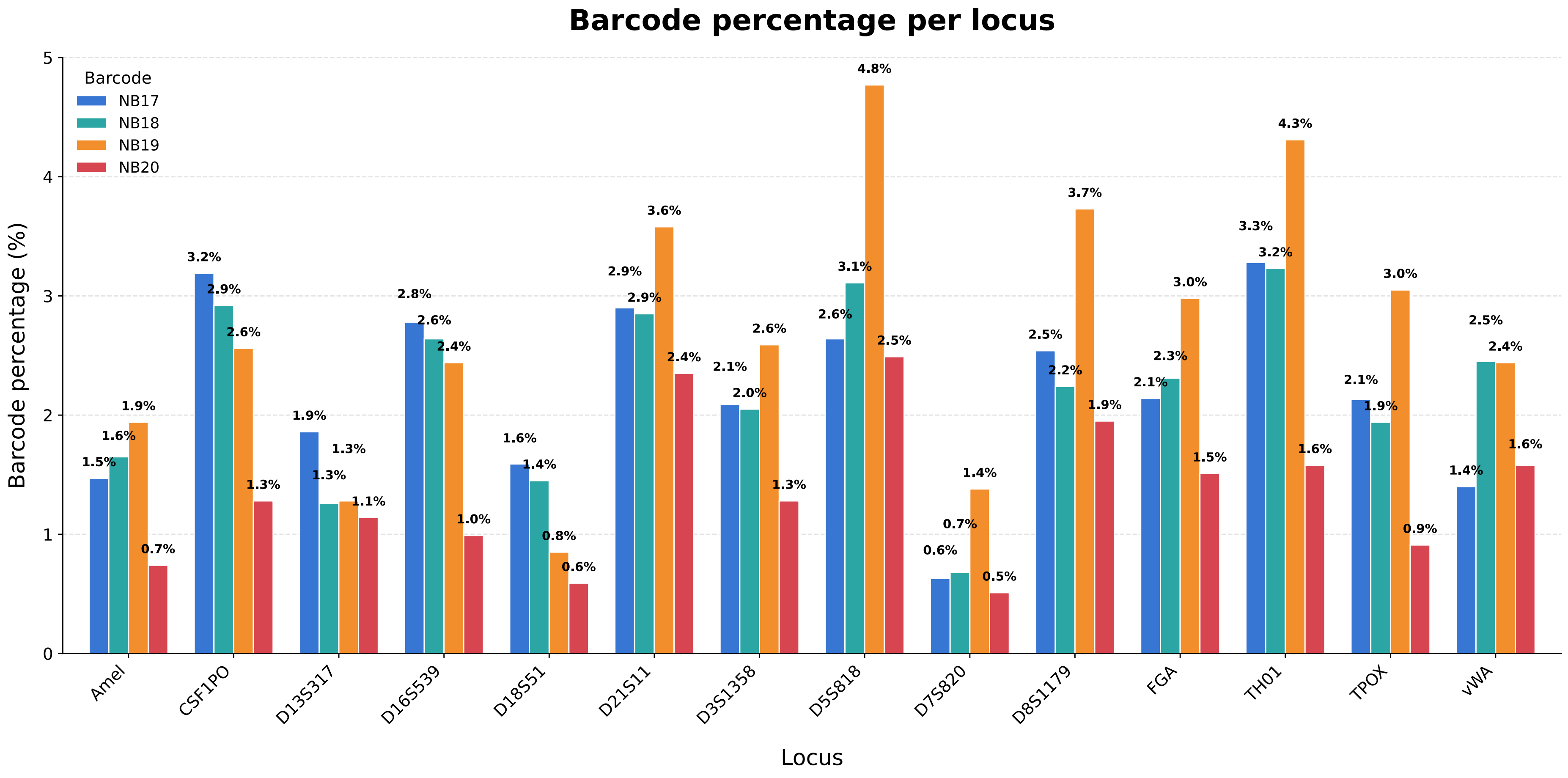

**Figure S4:** Readsaber plots of sample 9947A and 9948 after RPA amplification at 34°C.
Forward and reverse primers are indicated by locusname + F or R, respectively. The target is indicated in grey and labelled as 'target'. Grey blocks labelled as 'N' are sequences of 10 bp or longer without annotations. The relative read count per annotation pattern (%) is given at the left side of the annotation patterns. The arrows indicate the orientation of the primer and target sequences.

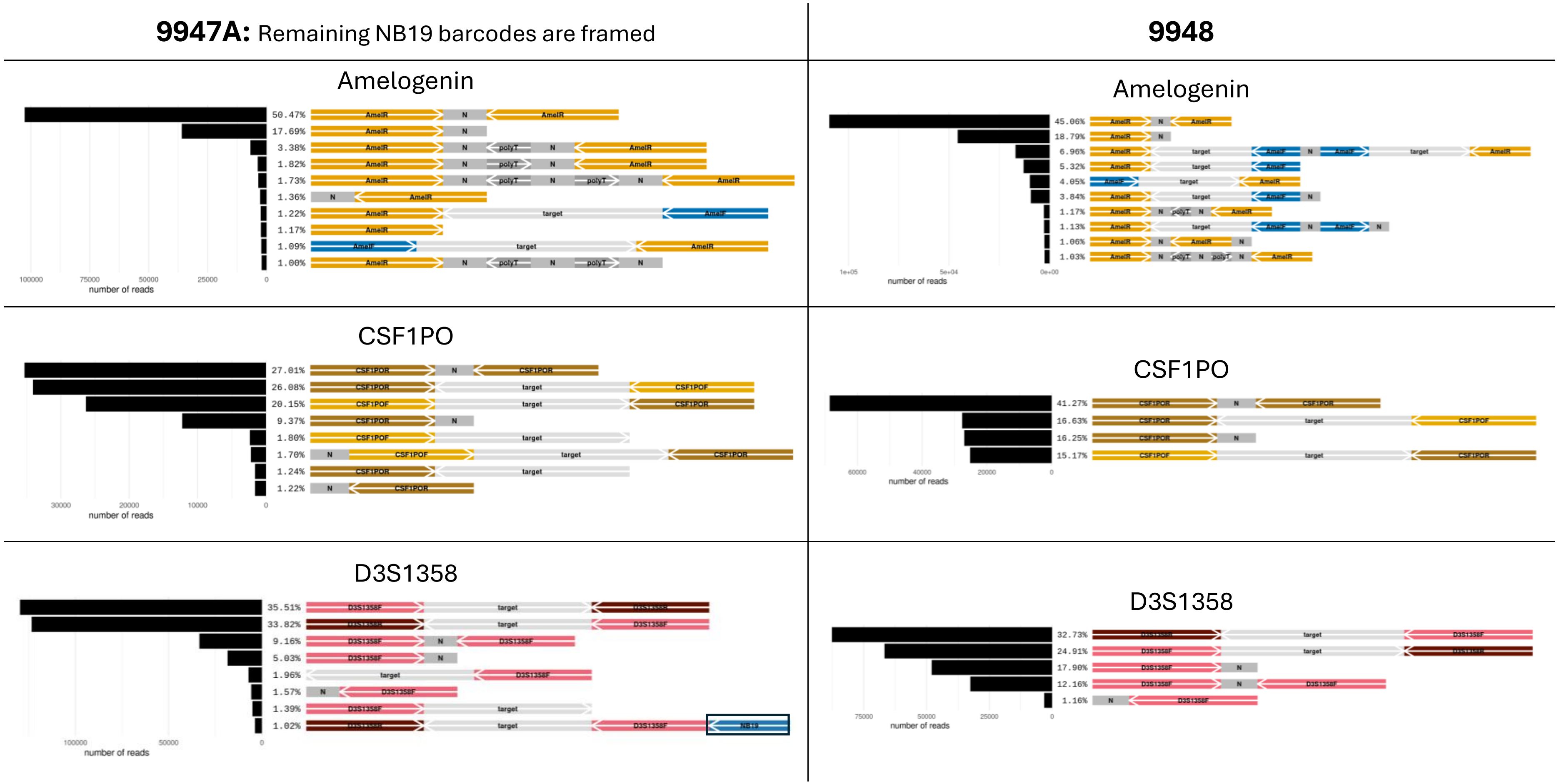

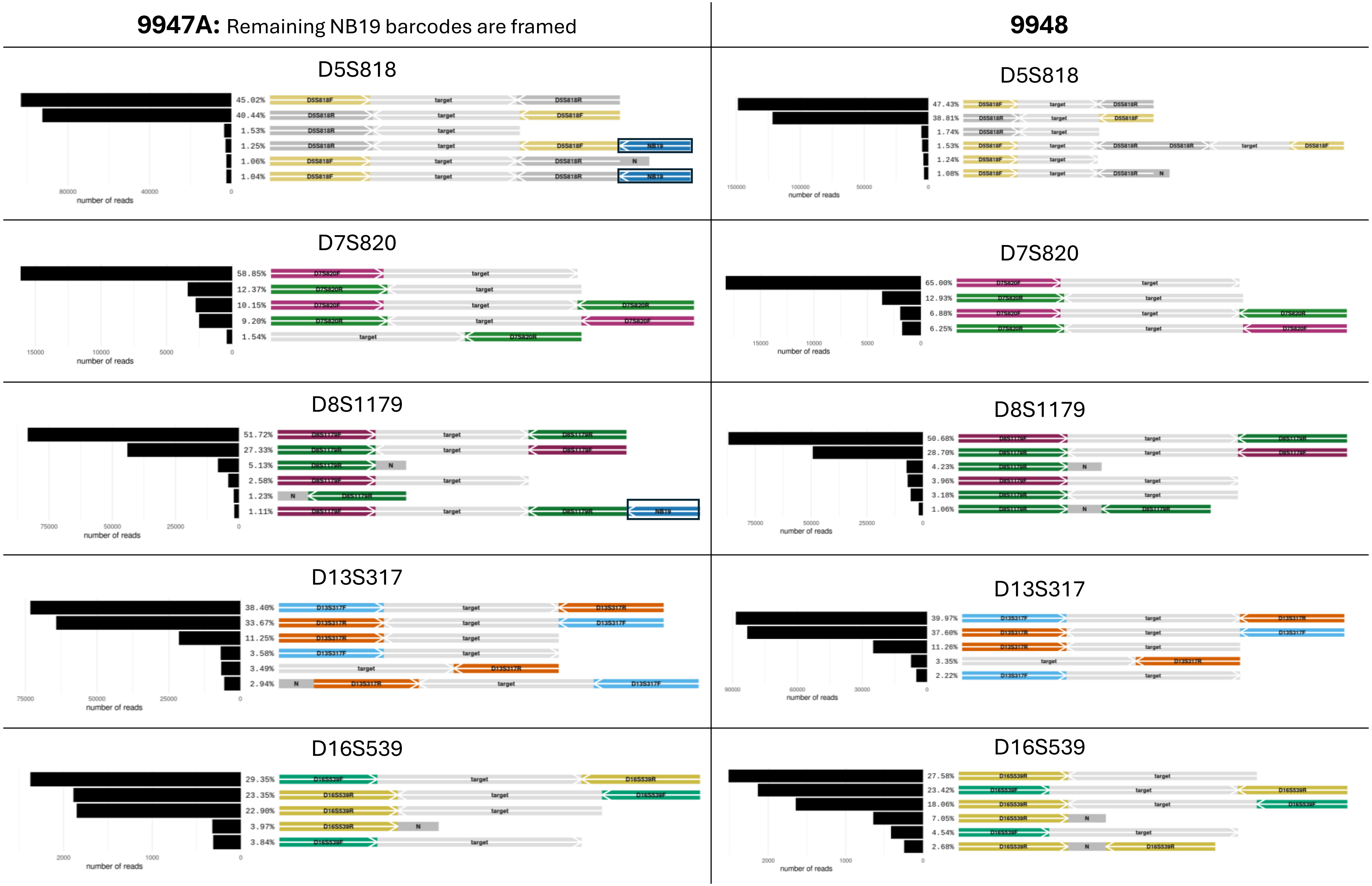
9947A: Remaining NB19 barcodes are framed
9948
D5S818
45.02% 40.44% 1.53% 1.25% 1.06% 1.04%
47.43% 38.81% 1.74% 1.53% 1.24% 1.08%
D7S820
58.85% 12.37% 10.15% 9.20% 1.54%
65.00% 12.93% 6.88% 6.25%
D8S1179
51.72% 27.33% 5.13% 2.58% 1.23% 1.11%
50.68% 28.70% 4.23% 3.96% 3.18% 1.06%
D13S317
38.40% 33.67% 11.25% 3.58% 3.49% 2.94%
39.97% 37.60% 11.26% 3.35% 2.22%
D16S539
29.35% 23.35% 22.90% 3.97% 3.84%
27.58% 23.42% 18.06% 7.05% 4.54% 2.68%
number of reads
target
N
NB19

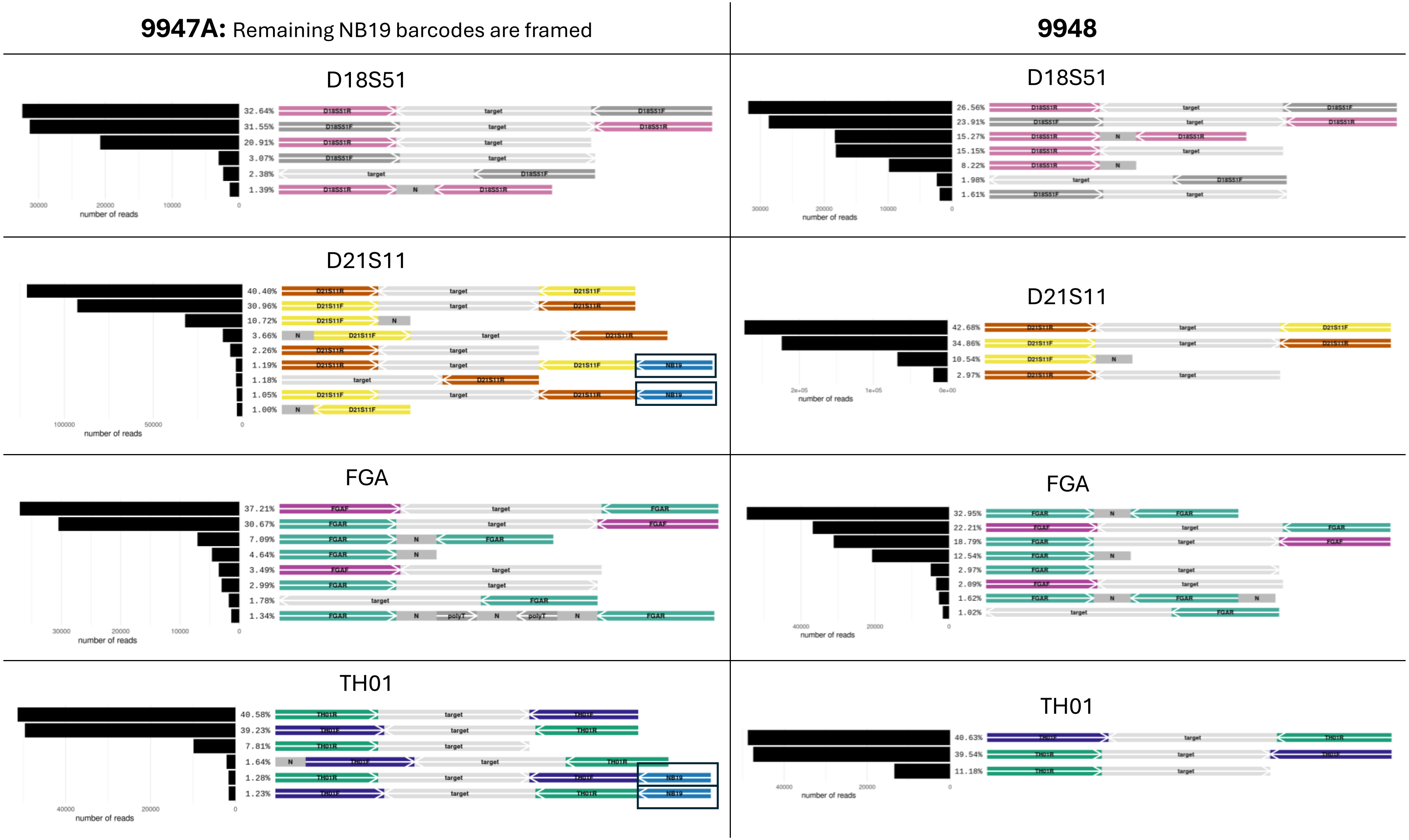
9947A: Remaining NB19 barcodes are framed
9948
D18S51
32.64%
31.55%
20.91%
3.07%
2.38%
1.39%
number of reads
D18S51
26.56%
23.91%
15.27%
15.15%
8.22%
1.98%
1.61%
number of reads
D21S11
40.40%
30.96%
10.72%
3.66%
2.26%
1.19%
1.18%
1.05%
1.00%
number of reads
D21S11
42.68%
34.86%
10.54%
2.97%
number of reads
FGA
37.21%
30.67%
7.09%
4.64%
3.49%
2.99%
1.78%
1.34%
number of reads
FGA
32.95%
22.21%
18.79%
12.54%
2.97%
2.09%
1.62%
1.02%
number of reads
TH01
40.58%
39.23%
7.81%
1.64%
1.28%
1.23%
number of reads
TH01
40.63%
39.54%
11.18%
number of reads

**9947A:** Remaining NB19 barcodes are framed

**9948**

## 9947A

##### TPOX

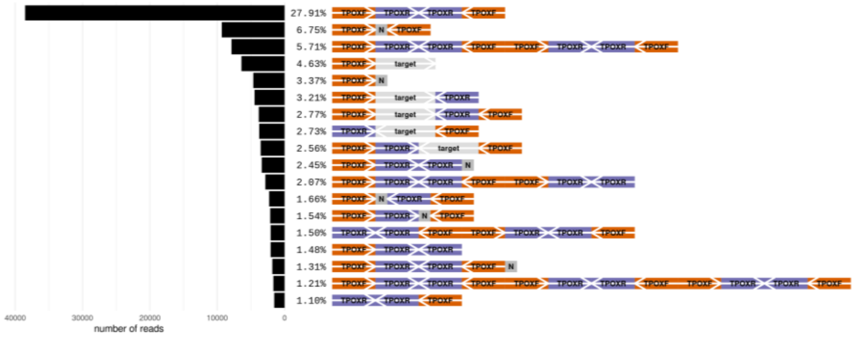


##### vWA

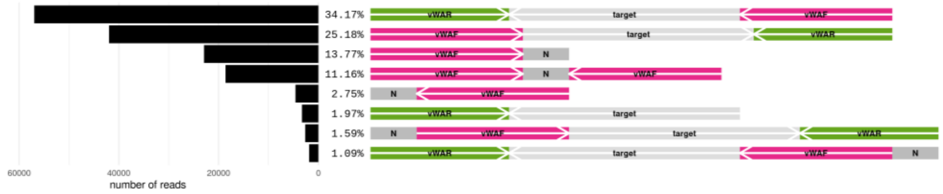


## 9948

##### TPOX

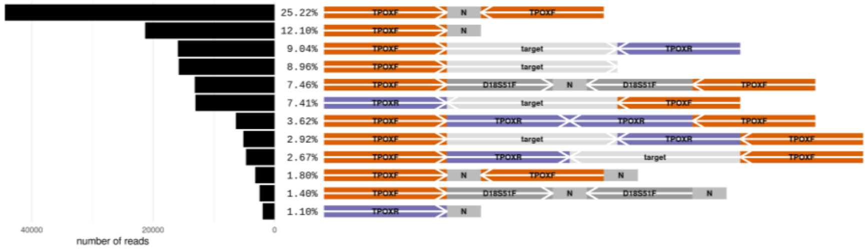


##### vWA

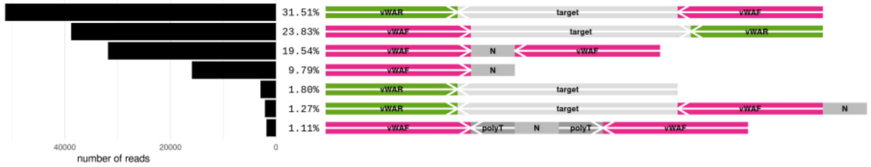

**Figure S5:** Readsaber plots of PCR data for 9947A, originating from data published by Tytgat et al. (2022). Forward and reverse primers are indicated by locusname + F or R, respectively. The target is indicated in grey and labelled as 'target'. Grey blocks labelled as 'N' are sequences of 10 bp or longer without annotations. 'N' sequences flanking the primers are the result of the ForenSeq DNA preparation, as Nanopore sequencing was in this study proceeded by Illumina sequencing. Because the indexes of this sample are unknown, these features could not be added to the annotation file. The relative read count per annotation pattern (%) is given at the left side of the annotation patterns. The arrows indicate the orientation of the primer and target sequences.

**9947A** (Referred to as 'sample f' by Tytgat et al.)

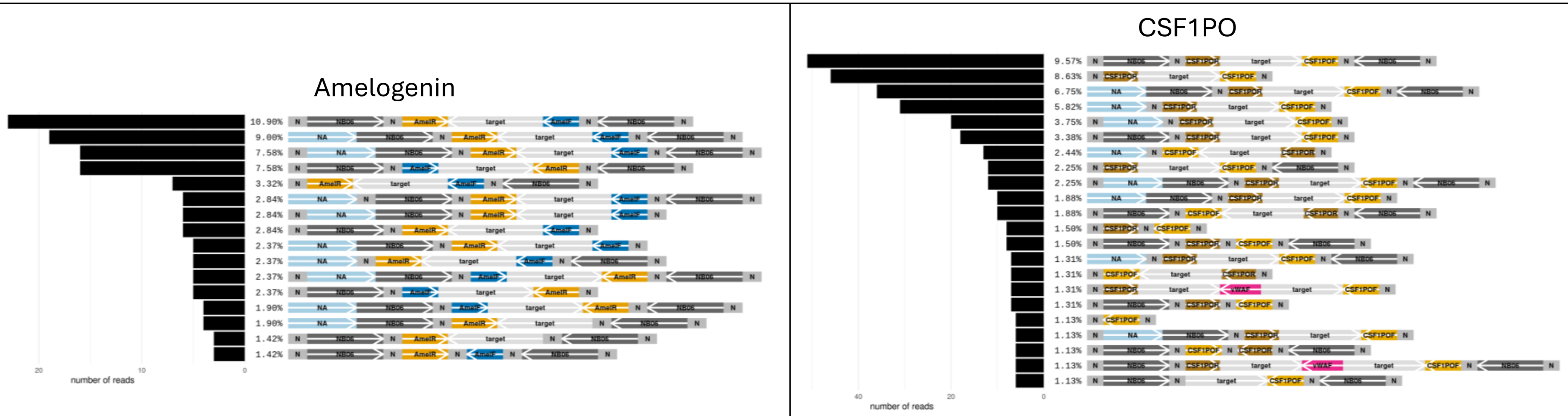

#### **9947A** (Referred to as 'sample f' by Tytgat et al.)

D3S1358

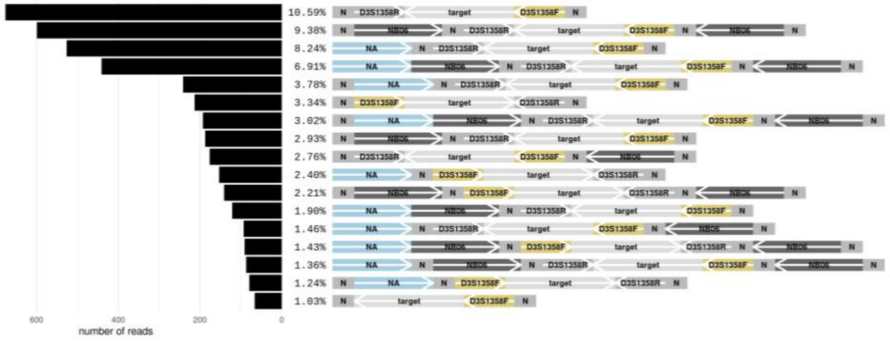


D5S818

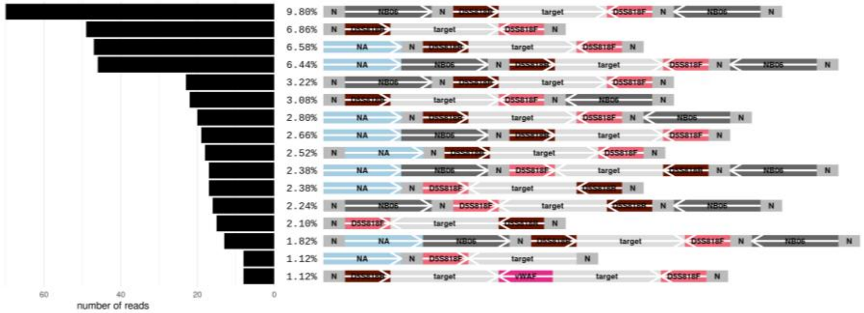


D7S820

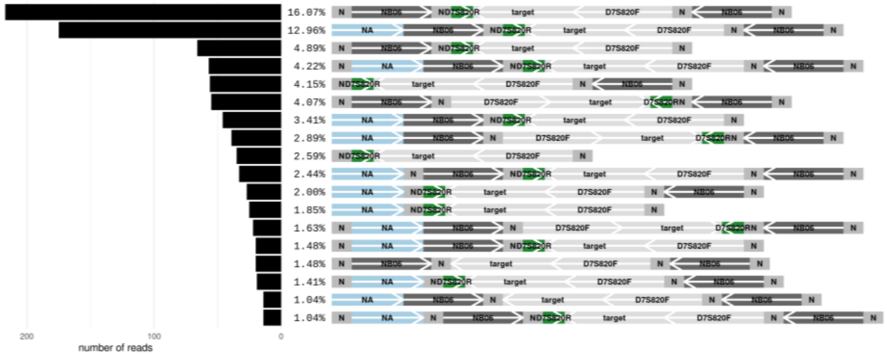


D8S1179

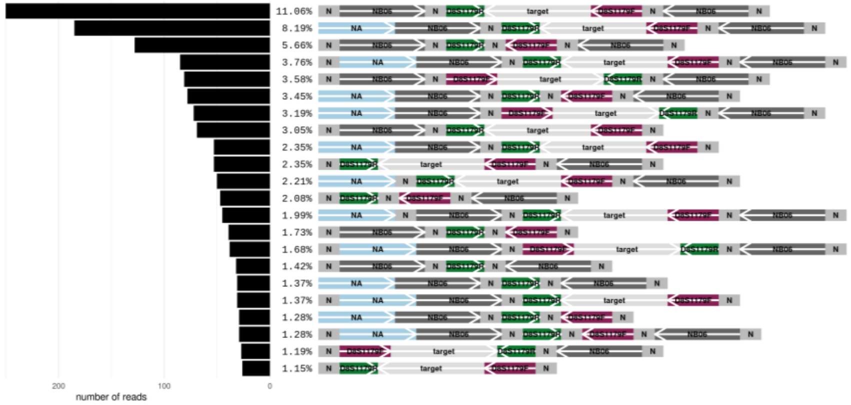

#### **9947A** (Referred to as 'sample f' by Tytgat et al.)

### D13S317

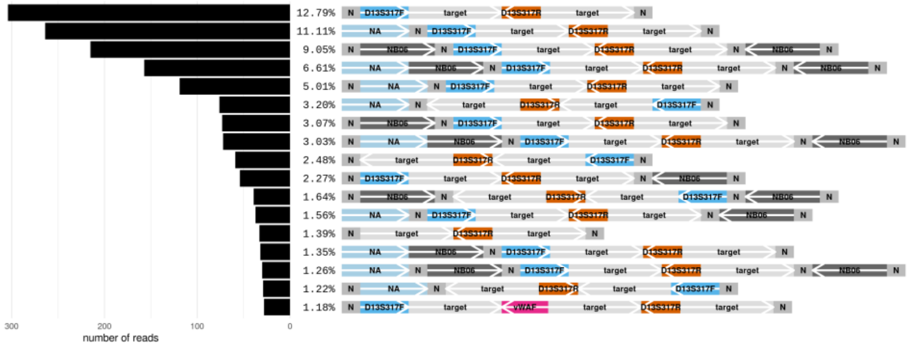


### D16S539

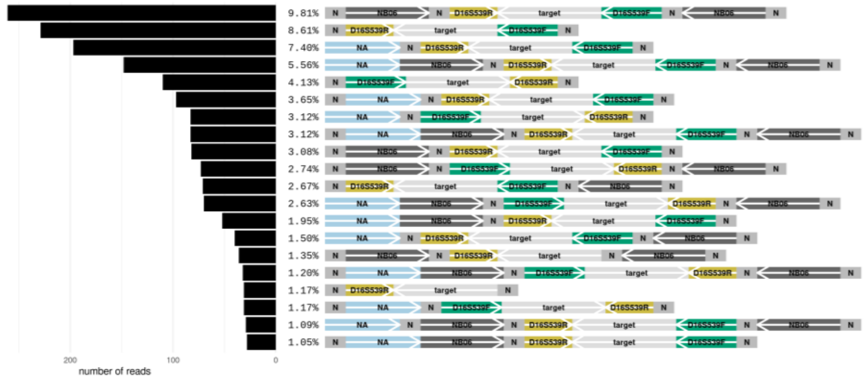


### D18S51

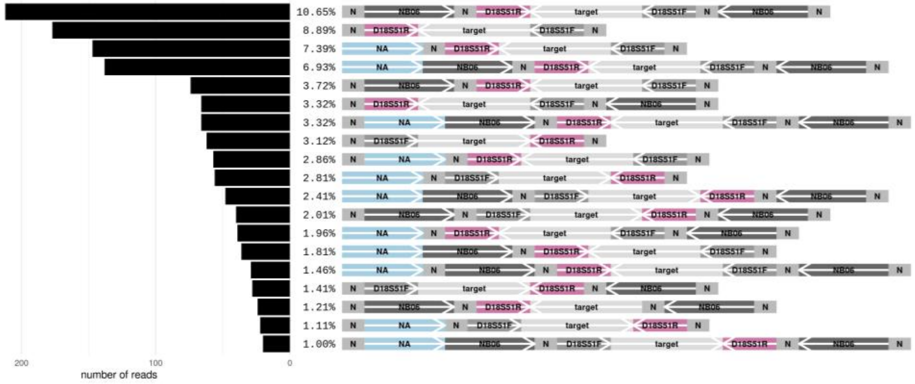


### D21S11

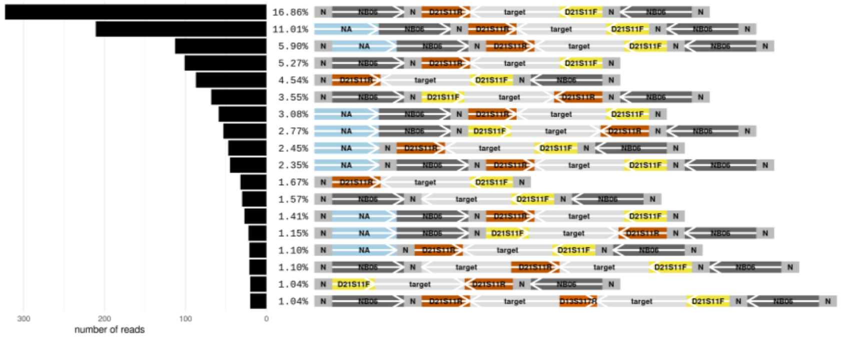

#### **9947A** (Referred to as ‘sample f’ by Tytgat et al.)

##### FGA

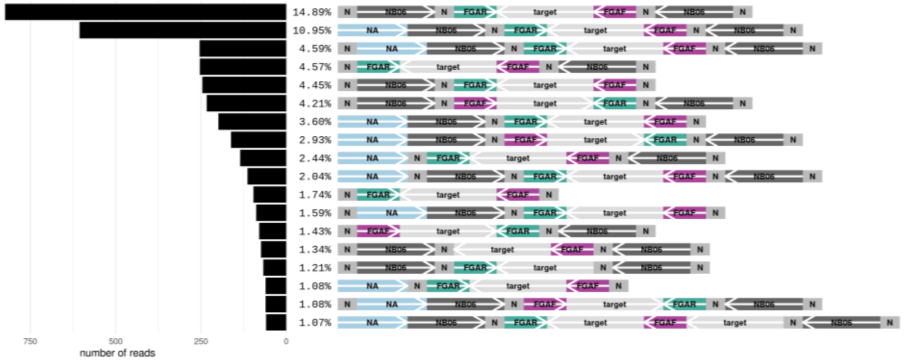


### TH01

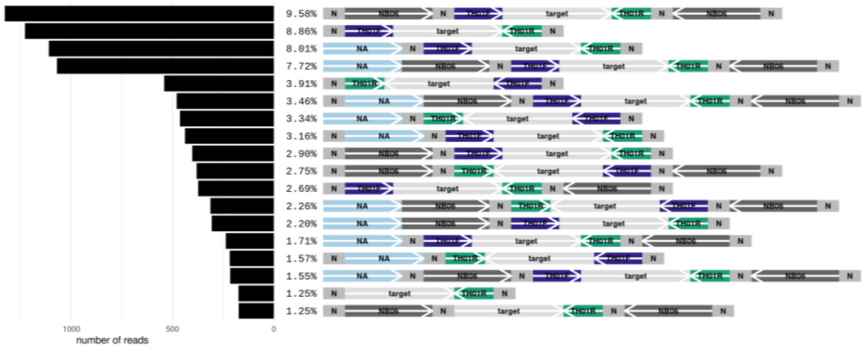


##### TPOX

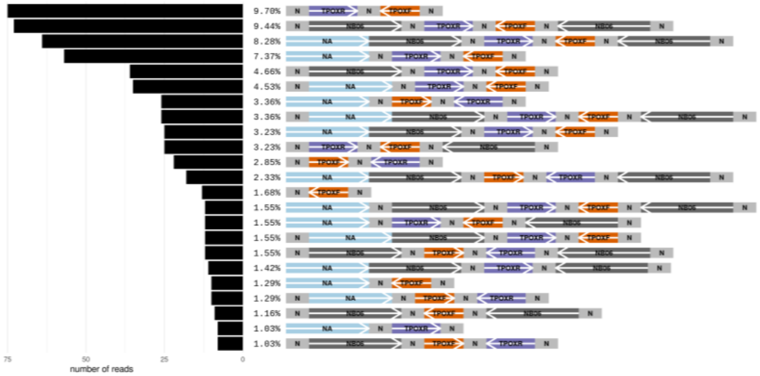


##### vWA

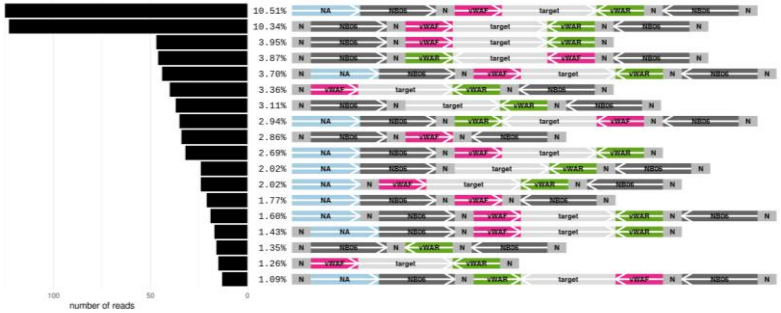

**Figure S6:** Electropherograms of the no-template controls (NTC) of sample 3.
The left and right panels show the NTCs for loci D3S1358 and D5S818, respectively. Peaks coloured in orange are separated from each other by multiples of the primer length, and where therefore classified as primer multimer peaks.

D3S1358

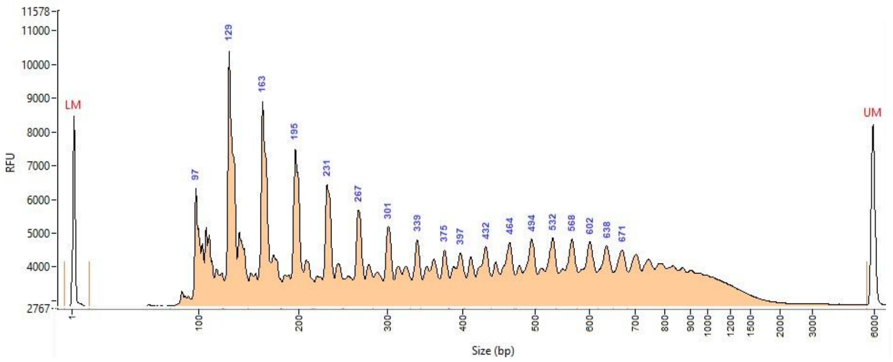


D5S818

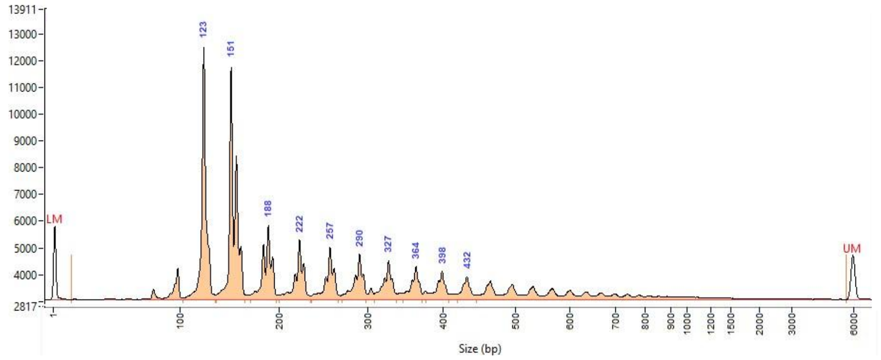
